# Larger larval sea lamprey (*Petromyzon marinus)* have longer survival times when exposed to the lampricide 3-trifluoromethyl-4-nitrophenol

**DOI:** 10.1101/2025.01.10.632359

**Authors:** Allison M. Nalesnik, Emily L. Martin, Ian S. Kovacs, Connor S. Johnson, Emma I. Carroll, Aaron Jubar, William Hemstrom, Michael P. Wilkie, Erin S. Dunlop, Maria S. Sepulveda, Nicholas S. Johnson, Mark R. Christie

## Abstract

Invasive sea lamprey (*Petromyzon marinus*) in the Laurentian Great Lakes have negatively impacted ecological and economically important fishes for nearly a century. To mitigate these effects, the lampricide 3-trifluoromethyl-4-nitrophenol (TFM) is applied annually on a rotating basis to selected Great Lakes tributaries to kill larval lamprey before they become juveniles, out-migrate to the lakes, and parasitize other fishes. It has been hypothesized that larval size (e.g., mass, length) may affect survival time in response to TFM. To test this hypothesis, we conducted an experiment with 8,611 larvae across four temporal replicates, in which TFM concentrations equivalent to those used in present-day stream treatments were applied for up to 18 hours. When examining the survival times of larval lamprey exposed to TFM, we found a significant, positive relationship between length, mass, toxicity, and their interactions. For every one mm increase in total length, a corresponding increase by one gram of mass reduced survival time by 0.4315 minutes [95% CI: 0.5283 – 0.2992] and vice versa (i.e., the significant interaction between length and mass revealed that as larvae increase in mass, the survival benefit to being longer decreases, and vice versa). The changes in total length and mass of larval sea lamprey stored in ethanol for four months was also quantified. The observation that five larvae survived well past the 12-hour time window of a typical TFM field treatment highlights the need for continuous monitoring and the development of new control strategies to ensure the continued effective management of this invasive species.

## Introduction

Sea lamprey (*Petromyzon marinus*) are hematophagous, non-host specific parasites that are native to the North Atlantic Ocean with populations along the coasts of Europe and North America. However, sea lamprey are an invasive pest in the Laurentian Great Lakes (Hansen et al., 2016) where they have contributed to substantial mortality and population declines in ecologically and economically important fishes (Coble et al., 1990; Kitchell, 1990). Sea lamprey were first documented in Lake Ontario in the late 1800s, after which they rapidly spread to Lake Erie following modifications to the Welland Canal in the early 1900s (Docker et al., 2021; Eshenroder 2014). By the late 1930s, sea lamprey had spread to Lake Huron, Lake Michigan, and Lake Superior (Christie and Goddard, 2003). In the novel environment of the Great Lakes, the typically anadromous sea lamprey acclimated to life-long residence in an entirely freshwater environment and developed smaller body sizes both at the time of metamorphosis and as adults in comparison to individuals from their native range (Hansen et al., 2016). These changes may be, at least in part, the result of a response to selection in genes responsible for growth, reproduction, and bioenergetics (Yin et al., 2021). This capacity for adaptation to novel environmental conditions suggests that sea lamprey may also be capable of responding to other selective pressures encountered in the Great Lakes.

Early control methods were initiated in the 1950s and relied mostly on engineered structures such as physical and electrical barriers and physical traps to prevent adult sea lamprey from accessing reproductive habitat (Hrodey et al., 2021; Smith and Tibbles, 1980), but the devices were labor intensive to operate and blocked valued fishes (McLaughlin et al., 2007). Lampricides were developed as an alternative approach to population control; over 4,000 organic candidate chemicals were tested at the Hammond Bay Biological Station (HBBS, Millersburg, MI), of which ten compounds were found to have greater toxicity for larval sea lamprey than other fishes at comparatively lower concentrations (Applegate et al., 1957, 1958, 1966). The chemical compound 3-trifluoromethyl-4-nitrophenol (hereafter TFM) was identified as the most effective agent and was found to be practical for field application, bearing relatively low toxicity to other species at low concentrations and a low manufacturing cost (Applegate et al., 1961; Smith and Tibbles, 1980).

TFM induces mortality by interfering with the oxygen-dependent production of ATP within cells and tissues (Birceanu et al. 2021; Huerta et al. 2020; Niblett and Ballantyne 1976). Larval sea lamprey are hypothesized to be particularly susceptible to TFM given their basal location on the vertebrate phylogeny and limited ability to metabolize TFM through its molecular conjugation to form TFM glucuronide, a less toxic TFM derivative that is readily excreted (Bussy et al., 2018; Kane et al. 1994; Lawrence et al., 2022; Lech and Statham, 1975). Since control methods were implemented in the late 1950s, sea lamprey populations have decreased by approximately 90% from their historically high abundances, aiding the recovery of some native fish populations such as lake trout (*Salvelinus namaycush;* Marsden and Siefkes, 2019). The continued control of sea lamprey in the Great Lakes hinges on effective stream treatments, which can be influenced by local environmental conditions at treatment sites. Because TFM bioavailability is higher in low pH and low alkalinity waters, control personnel can use this relationship to modify TFM treatment concentrations accordingly (Wilkie et al., 2021). Larval TFM sensitivities may also be influenced by water temperature (Hlina et al., 2021; Muhametsafina et al., 2019), time of year (Scholefield et al., 2008), and possibly liver glycogen content (although direct mechanisms are not well described), such that treatment efficacy can depend on water conditions and larval body conditions (Schueller et al., 2024). Despite TFM being instrumental to controlling sea lamprey populations, specific quantitative relationships between larval lamprey body size and toxicant susceptibility remain largely uncharacterized, although TFM uptake is known to be inversely related to body mass (Tessier et al., 2018). In the present study, we examine how total length, mass, and their interaction contribute to TFM susceptibility in sea lamprey. A better understanding of these effects is critical for determining appropriate TFM dosages and durations for field applications over the next few decades, which can be adjusted based on larval size surveys to ensure continued treatment effectiveness and population control.

Here, we empirically test the relationship between larval sea lamprey body size and survival under TFM exposure while accounting for pH, season, temperature, and source of larvae. We predicted that because oxygen consumption and TFM uptake per unit of mass are greater in smaller mass larvae (Tessier et al., 2018), larger animals would be more likely to survive for prolonged periods of time in TFM. To test this hypothesis, we exposed 8,611 larval sea lamprey to TFM concentrations designed to mimic treatments in the field and measured survival after each hour of exposure. We measured the total length and mass of each larva and quantified the relationships between these morphometric measures and survival time. We also used this opportunity to describe the relationship between changes in body size in larval sea lamprey and their storage time in ethanol. To do so, we quantified changes in total length, mass, and condition of larval sea lamprey over a four-month period.

## Materials and Methods

### Sample collection

All larvae used in this experiment were collected by crews from the United States Fish and Wildlife Service and United States Geological Survey over a period of eight days during July 2022 (ESM **Table S1**). Larvae were obtained from the Muskegon River, a tributary to Lake Michigan, MI, as it harbors one of the largest populations of spawning adult and larval sea lamprey within the Great Lakes basin (Lavis et al., 2003; Slade et al., 2003). Prior to collecting the larval lamprey for the present study, the Muskegon River had been treated with TFM in 2017 and 2019. Given this three-year interval and the growth rate of larvae (Dawson et al., 2021), the larval lamprey used in this study are unlikely to have been previously exposed to TFM. Pulsed-DC backpack electrofishing units were used to capture larval sea lamprey from four reaches of the Muskegon River (ESM **Table S1**). To minimize the effects of seasonality, we condensed collection times as much as possible. Most larvae collected were between 60-100 mm (range: 40-140 mm).

To minimize stress experienced by the larvae due to handling and transportation, larvae were transferred to acclimation tanks at U.S. Geological Survey, Hammond Bay Biological Station (HBBS; Millersburg, MI) within one day of collection in individually labeled zippered mesh bags (Tutudow, 40 cm x 50 cm, polyester, 250 µm mesh). Up to 50 larvae were transported in each bag alongside two liters of sand from the Muskegon River. The sand grains were larger than the mesh size of the bag such that the sand added to each bag remained in place during transit. Bags containing larvae and sand were placed in 50 L aerated tanks maintained at a temperature of 15 °C overnight (approximately the temperature of the Muskegon River). The day after capture, larvae held in mesh bags were transported to HBBS in 200 L aerated tanks maintained at 15 °C.

Upon arrival at HBBS, larvae in each bag were separated from the sand by emptying the contents of each bag onto a 750 µm screen and using a slow stream of water from Lake Huron at 15 °C to wash the sand through the screen, leaving the larvae on top of the screen. Larvae from each specific bag were next moved into a mesh container (Ikea Fyllen Laundry Basket, 79 L volume, 50 cm high and 45 cm diameter, 500 µm mesh size) and placed inside an aerated 1000 L tank supplied with water from Lake Huron at ambient temp (16-18 °C; water was pumped from a depth of 25 m in Lake Huron). Within each holding tank, six mesh containers were used to hold at most 282 larvae each. Sand with a 250 µm grain size was added to each mesh container to achieve a layer of sand five cm thick for the larval lamprey to burrow into. Capture dates and locations, the dates of transportation, and the locations of acclimation at HBBS were tracked for each batch of larvae (ESM **Table S1**). All larvae were acclimated at HBBS within the mesh container they were assigned for two weeks prior to exposure to TFM, during which larvae were fed fresh baker’s dry yeast at a rate of 1 gram (g) of yeast per larvae two times per week (Hanson et al., 1974; Hume et al., 2024).

### TFM exposure experiments

Our TFM target concentrations were determined by preliminary range-finding trials to determine the lethal concentration of TFM required to kill 90% of larval sea lamprey over 12 hours (12-h LC_90_), based on the local water conditions and the tank systems to be used, following the standard guidelines for acute toxicity tests in fish (OECD, 2019). All protocols involving the handling of larvae at the Hammond Bay Biological Station were carried out in accordance with United States federal guidelines for care and use of animals in accordance with American Fisheries Society, ARRIVE guidelines, and the U.S. Animal Welfare Act of 1970 (Jenkins et al., 2014). We aimed to use a 12-hour exposure period to approximate the total duration of TFM treatments in the field (Middaugh et al., 2014). Our study consisted of four replicate TFM exposure experiments (replicated two days apart) each using an identical design to measure the survival of larval sea lamprey exposed to TFM. Each replicate experiment was conducted in two 1500 L tanks containing 624 L of water supplied from Lake Huron. Each tank was dosed with between 104 and 112 mL of a 15,000 mg/L TFM stock solution at least 12 hours prior to introducing the larvae to reach target concentrations of 2.3 to 2.5 mg/L TFM. No additional water was added to these tanks after TFM was added. Water from Lake Huron did not vary substantially in temperature (μ = mean ± standard deviation; μ = 17.6 ± 0.5 °C) or alkalinity over the four replicant trials (μ = 85.6 ± 0.6 mg/L CaCO_3_; ESM **Table S2**). However, in some cases, there were fluctuations in pH during treatments, as discussed below (μ = 8.1 ± 0.3; ESM **Table S2**). Each tank was fitted with a 15 cm long aeration stone at the center to facilitate uniform distribution of TFM throughout the tank and to keep dissolved oxygen (DO) concentrations higher than 7.6 mg/L (μ = 9.0 ± 0.6 mg/L; ESM **Table S2**). The air stone was set to turbulently aerate the water for at least 30 minutes after initial dosing and reduced to a moderate level for subsequent mixing and aeration. The TFM concentration was measured thirty minutes after initial dosing the day before each experiment began and at least 12 hours later on the day of the experiment to determine if additional adjustments were required to maintain consistent exposure conditions between our replicates.

Between 55 and 282 larval sea lamprey were kept in each of six mesh bags during acclimation (ESM **Table S1**) which were transferred to the experimental tank at the start of each experiment. When the bags in the acclimation tanks were lifted and twisted, the 250µm sand fell through the 500µm mesh containers leaving only larval sea lamprey and no sand, eliminating the stress and potential injuries to larvae if each larva was dug from the sand. The larvae in each bag were then lowered into the experimental tank nearby. Each bag was assessed for mortality in the order in which the bags went into the experimental tanks.

Throughout the duration of each experiment, several parameters were monitored on an hourly basis. Fish mortality within each bag was assessed to determine survival time in TFM. To assess mortality, teams of two people transferred the larvae from one mesh bag into a double-net apparatus (**SI video**). Fish that were alive with swimming ability intact swam through the apparatus and into an empty mesh bag held below. Dead or impaired fish were unable to swim through the apparatus and required visual inspection to determine status (alive or dead). Larvae were inspected in a shallow plastic tray filled with the same TFM solution from the respective experimental tank for up to three minutes. Larvae were declared dead after the cessation of branchiopore (gill pore) movement and unresponsiveness to tail pinching, then stored in individually labeled bottles of 95% ethanol. Impaired fish were returned to their original experiment tank and into the same mesh bag as individuals that swam through the apparatus. In addition to recording larval sea lamprey mortality at hourly intervals, DO (YSI Pro 20), pH (Fisherbrand Accumet AP110 Portable), salinity (YSI Pro 20), TFM concentration, water and air temperature were measured hourly. TFM concentration was measured using a Genesys 6 spectrophotometer (Thermo Electron Corporation, MA, USA) set to a wavelength of 395, using analytical standards prepared in the same water, based on Standard Operating Procedure No. LAB 431.0, Hammond Bay Biological Station, Millersburg, MI (GLFC. https://glfc.org/sop.php). Analytical standards were prepared from TFM (CAS Number 88-30-2; Millipore-Sigma, St. Louis, MO). Water and air temperature were measured via a laboratory liquid-in-glass thermometer. The experiments varied in duration from five to eighteen hours until the last larva had died. At the conclusion of all experiments, preserved larvae were transported from HBBS to Purdue University (West Lafayette, IN) where the total length (mm) and mass (g) was measured for all lamprey (n = 8,611 larvae). All experiment samples were stored in 95% ethanol for at least four months prior to measuring their total length and mass.

To determine what physiological variables impact lamprey survival time in TFM, we used model selection to identify the best fitting linear mixed effect models for predicting survival time (hour), using total length (mm), mass (g), toxicity units, and their interactions. Toxicity units were calculated by the Henderson-Hasselbalch equation (McDonald & Kolar 2007) as 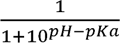 where the *pKa* (the negative log of the acid dissociation constant, Ka, for an acid-base reaction) is 6.07 for TFM (Hubert 2003). These variables were fitted as fixed effects; we also included a nested random effect of experiment in tank in bag as a covariate using R 4.3.2 (R Core Team, 2023) and the R package lmerTest (Kuznetsova et al., 2017). Based on diagnostic assessments, the assumptions for a linear mixed effects model were largely met. A Q-Q plot of the residuals indicated that the data follow a normal distribution, with only minor deviations observed. Additionally, the residuals versus fitted values plot showed no discernible patterns, with most points randomly scattered around the zero line. While five data points exhibited slight deviations from the expected trend, given the dataset’s size, their impact on the model was minimal. Therefore, the assumptions of normality, linearity, and homoscedasticity were reasonably satisfied. We compared models using total length, mass, toxicity units, and their interactions (seen ESM **Table S3** for all models). Additionally, one tank (S4) within replicate one had a large fluctuation in toxicity units (illustrated in ESM **Fig. S1**). We evaluated our linear mixed effect models with and without this replicate and the best model was robust to its removal.We removed tank S4 within replicate one from our analyses due to the greater variance in toxicity units that is dissimilar from other replicates.

### Long-term effects of ethanol storage

To assess the effect of long-term ethanol storage on larval sea lamprey morphometrics, we systematically measured the total length and mass of two different sets of larval sea lamprey over the course of four months (hereafter storage experiments A and B). None of the larvae used to quantify changes in body size and mass due to ethanol storage were included in the TFM exposure experiments; instead, these larvae were used to determine the effect of measuring the 8,611 experimental larvae after a storage period of up to eight months since it was logistically impractical to measure all individuals during the large TFM exposure experiments. Storage experiment A: In July, prior to the large TFM survival experiment described above, 554 larval sea lamprey were transported to Purdue University’s Aquaculture Research Laboratory (ARL; West Lafayette, IN). Sea lamprey were exposed to TFM concentrations ranging from 2.0 mg/L to 4.0 mg/L in groups ranging from 10 to 25 individuals. If sea lamprey remained alive beyond 36 hours of TFM exposure, they were euthanized with 250 mg/L tricaine methanesulfonate (MS-222) buffered to pH 7 with sodium bicarbonate. Total length (mm) and mass (g) were measured immediately after death and each sample was individually stored in a 50 mL conical tube containing 40 mL of 95% non-denatured ethanol. Purdue University Institutional Animal Care and Use Committee reviewed experimental and handling procedures for this study, but did not require animal care protocols for the larval sea lamprey. Storage experiment B: At HBBS, 110 larval sea lamprey that remained in the acclimation tank after the completion of the large TFM survival experiments were euthanized with MS-222 using the same procedure described above. We measured the total length (mm) and mass (g) of each larva immediately after death, then stored every individual in 15 mL conical tubes containing 12 mL of 95% ethanol. Thereafter, each larval sea lamprey (554 from ARL, 110 from HBBS) was measured for body mass and total length systematically over a period of 120 days (see **SI Methods** for details on measurement procedures). We calculated condition (CF) using the same equation as above.

Given that larvae become smaller over time when stored in ethanol, we fit an additional model to predict the initial size of an individual from a measurement taken after extended storage in ethanol. We first randomly sampled our dataset of Experiment A and Experiment B individuals (664 larvae) such that only one sample per individual was used (since any models containing individual ID as a covariate could not be used to make predictions for new individuals). We then used the *nls* function in R to fit a nonlinear regression model to determine the decay rate which best fit our observed data using an exponential decay function:

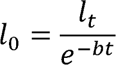

where *l_0_*is initial length, *l_t_* is the length at measurement time *t*, and *b* is the rate of decay. We evaluated the ability of this model to predict the initial size of a larval sea lamprey prior to storage using a hold-one-out cross validation approach (see **Tables S4** and **S5**; Wong, 2015). Using these adjusted values, we also re-analyzed the data from our large TFM exposure and body size experiments using the exact statistical models described above.

## Results

### Results from TFM exposure experiments

The 7,651 lamprey used in our four experimental replicates (seven experiment tanks) survived between two and 18 hours of TFM exposure (**Fig. 1A**). There was an allometric relationship between larval total length and larval mass (**Fig. 1B**). Measured TFM concentrations during exposure averaged 2.503 ± 0.07 mg/L (n = 78) across our replicates (**ESM Table S2; ESM Fig. S1**). TFM toxicity units averaged 0.03216, ranging from 0.009558 to 0.04813 (calculated from **ESM Table S2**). The 7,651 lamprey used in our four experimental replicates had total lengths between 43.0 and 137.5 mm (µ = 85.53 mm; **Fig. 1B**; **ESM Fig. S2A**) and masses between 0.074 and 4.529 g (µ = 0.8702 g; **Fig. 1C**; **ESM Fig. S2B**). The best fitting linear mixed effect model had the equation y= 6.711 + 0.02686*(total length, mm) + 0.4590*(mass, g) – 81.76*(toxicity units) – 0.006892(total length*mass) – 0.7365(total length*toxicity units) – 21.15(mass*toxicity units) and explained a large portion of the variation in survival time in TFM (conditional R² = 0.8790; **Table 1**; extended results in ESM **Table S3**). Total length was significantly and positively associated with survival time in TFM (**Table 1; ESM Fig. S2A)** (*p* value = 2.592•10^-8^). There was a 16.11 minute [95% CI: 10.46-21.80] increase in survival time for each one cm increase in total length after accounting for the effects of other parameters. Mass was significantly and positively associated with survival time in TFM (**Table 1**; ESM **Fig. S2B)** (*p* value = 0.01455). There was a 27.54 minute [95% CI: 5.504-49.68] increase in survival time for each one g increase in total mass after accounting for the effects of other parameters. Toxicity units were negatively associated with survival time, such that an increase in toxicity units leads to a decrease in survival time (*p* value = 1.180•10^-16^). There was a 98.11 minute reduction in survival time for each 0.02 increase in toxicity units. The effect of the interaction between total length and mass on survival time was negative (*p* value < 0.001): for every one mm increase in total length, a corresponding increase by one g of mass reduced survival time by 0.4315 minutes [95% CI: 0.5283 – 0.2992] and vice versa (*i.e.,* the significant interaction between total length and mass revealed that as larvae increase in mass, the survival benefit to being longer decreases, and vice versa) (**Fig. 2**; **Table 1**). The effect of the interaction between mass and toxicity units on survival time was positive (*p* value = 2.143•10^-7^): for every one g increase in mass, the impact of increasing by 0.02 toxicity units increases survival time by 25.38 minutes [95% CI: 15.80– 34.96]. The effect of the interaction between total length and toxicity units on survival time was negative (*p* value = 6.577•10^-7^): for every 10 mm increase in total length, the impact of increasing by 0.02 toxicity units reduces survival time by 8.838 minutes [95% CI: 12.32– 5.365]. Five individuals survived beyond 12 hours of TFM exposure (**Fig. 3; ESM Fig. S4**).

**Figure 1.**
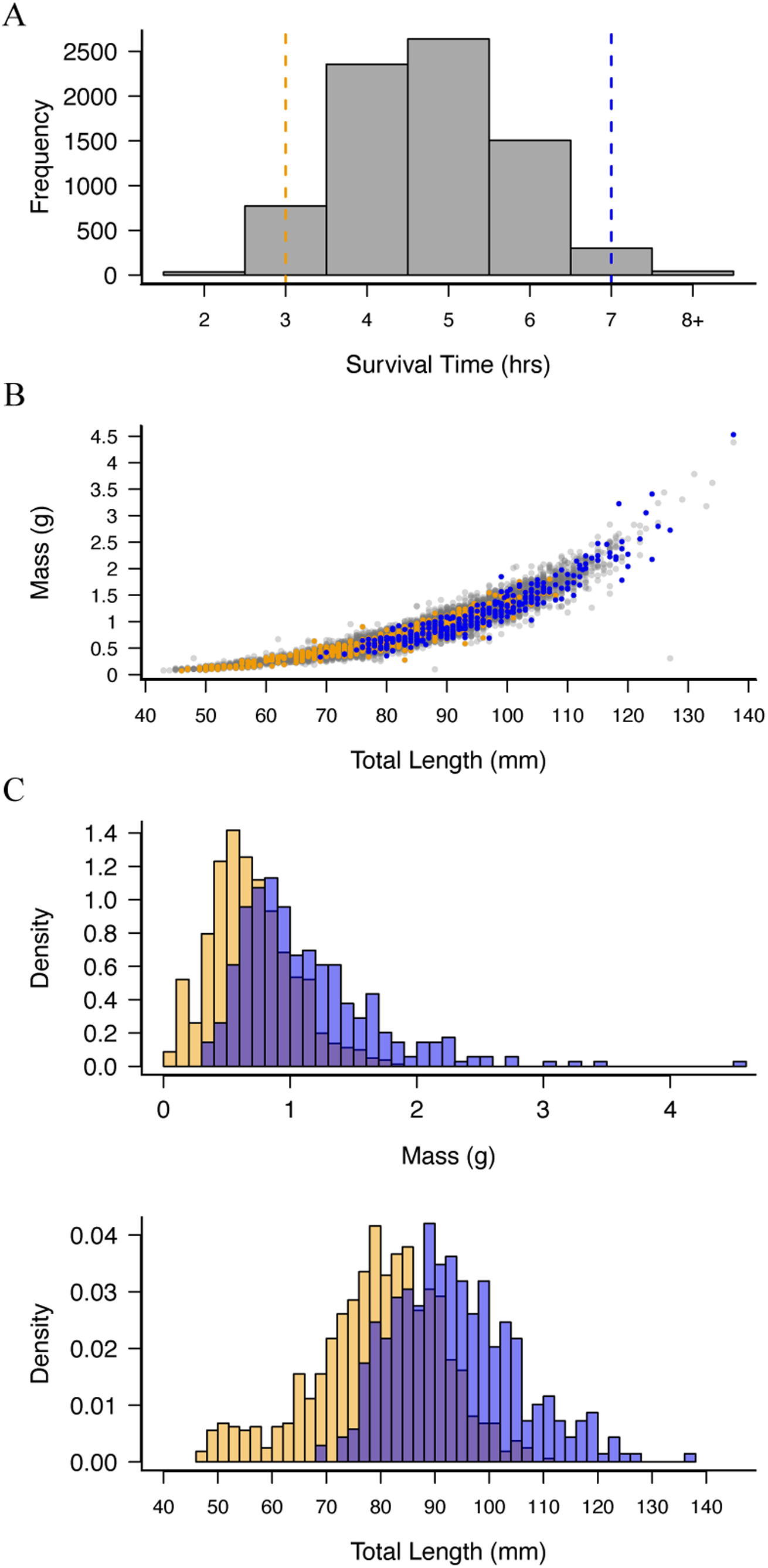
Distribution of sea lamprey exposed to TFM. The 7,651 lamprey exposed to TFM (2.52 ± 0.08 mg/L) survived between two and 18 hours of exposure (A). There was a highly allometric relationship between larval total length (mm) and larval mass (g), as illustrated by the 805 fish surviving less than three hours of exposure, representing 10.5% of our samples (orange; referred to as lower quantile) and the 345 fish surviving seven hours or beyond of exposure, representing the upper 4.51% of our samples (blue; referred to as upper quantile) (B). The lower quantile lamprey had masses between 0.074 and 1.801 g (µ = 0.694 g; orange bars; C top) and total lengths between 46.0 and 111.0 mm (µ = 80.18 mm; orange bars; C bottom). The upper quantile lamprey had masses between 0.333 and 4.529 g (µ = 1.129 g; blue bars; C top) and total lengths between 69.0 and 137.5 mm (µ = 94.27 mm; blue bars; C bottom).

**Figure 2.**
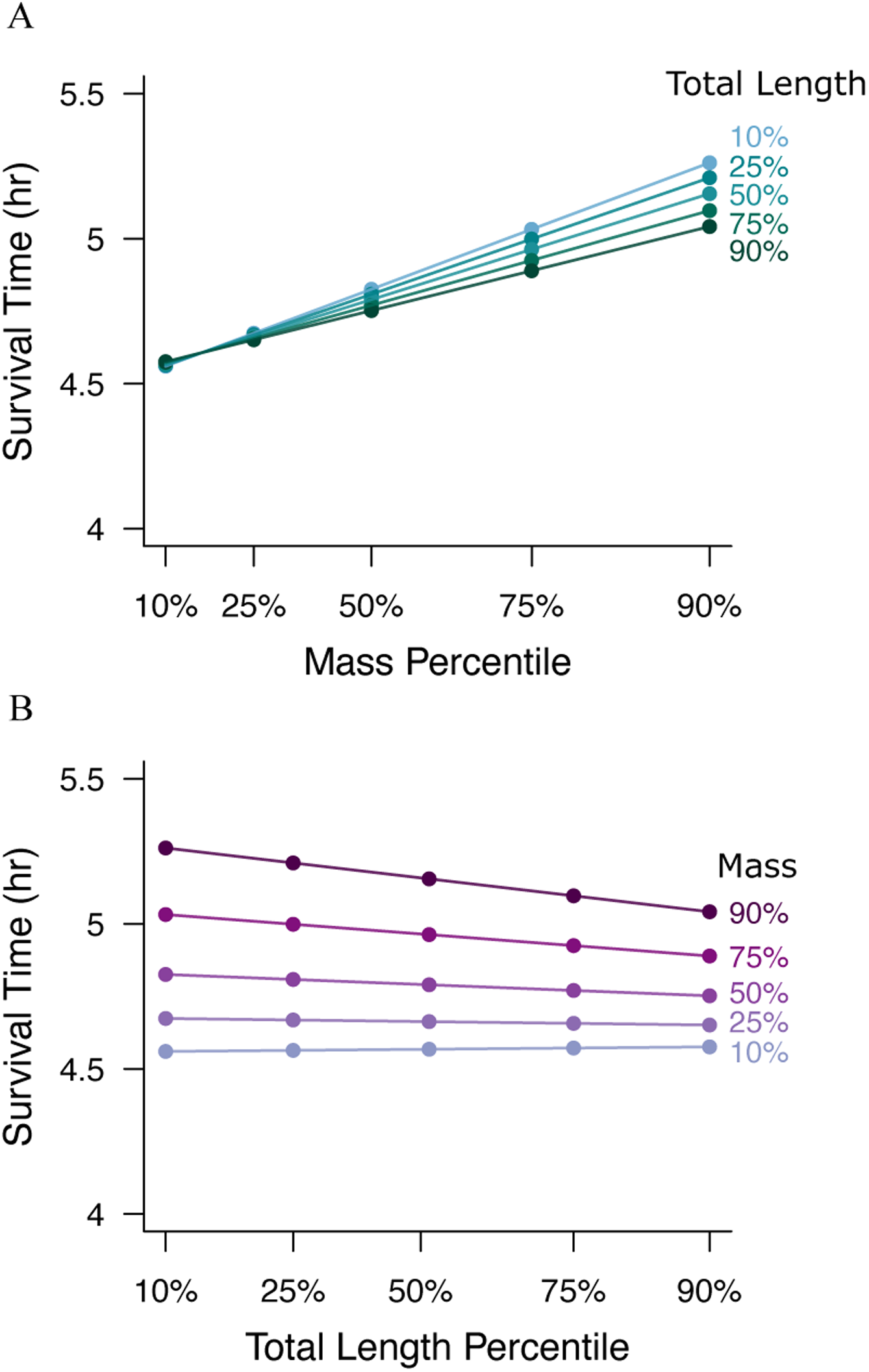
Predicted survival time (hrs) for the 10^th^, 25^th^, 50^th^, 75^th^, and 90^th^ mass and total length quantiles. **(A)** Relationship between mass (g) quantiles and predicted survival time (hrs) calculated using Model A (reported in text; see Table 1 for full model results). Line colors indicate the corresponding total length quantile. (B) Relationship between total length (mm) quantiles and predicted survival time (hrs) based on Model A (reported in text; see Table 1 for full model results). Line colors indicate the corresponding mass quantile.

**Figure 3.**
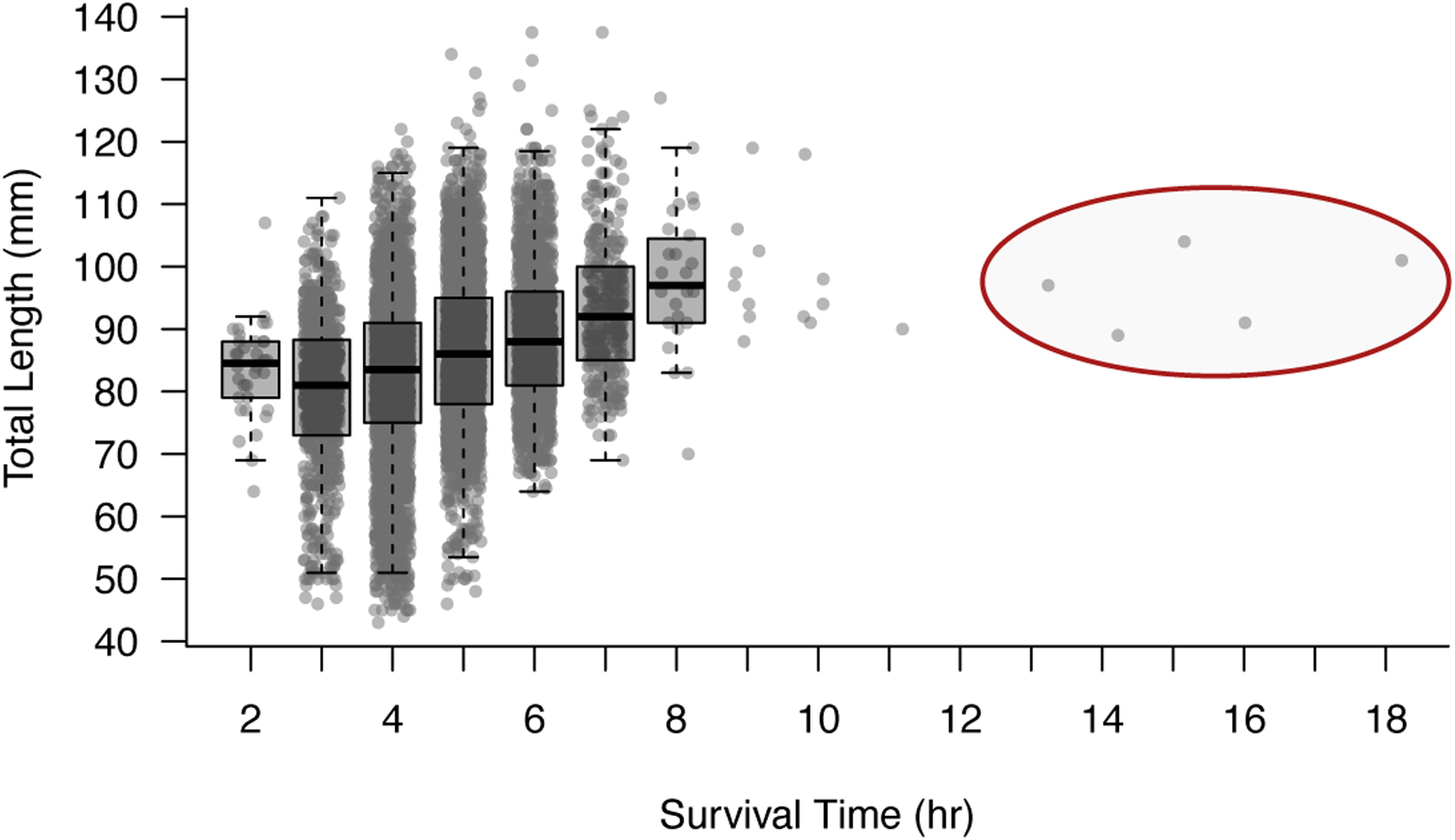
Relationship between body length and survival time of larval sea lamprey exposed to TFM (2.52 ± 0.08 mg/L) for up to 18 hours (n = 7,651 larvae). Zero mortality occurred at hours one and 12. Five larval sea lamprey survived beyond 12 hours of TFM exposure (circled in red).

**Table 1.**
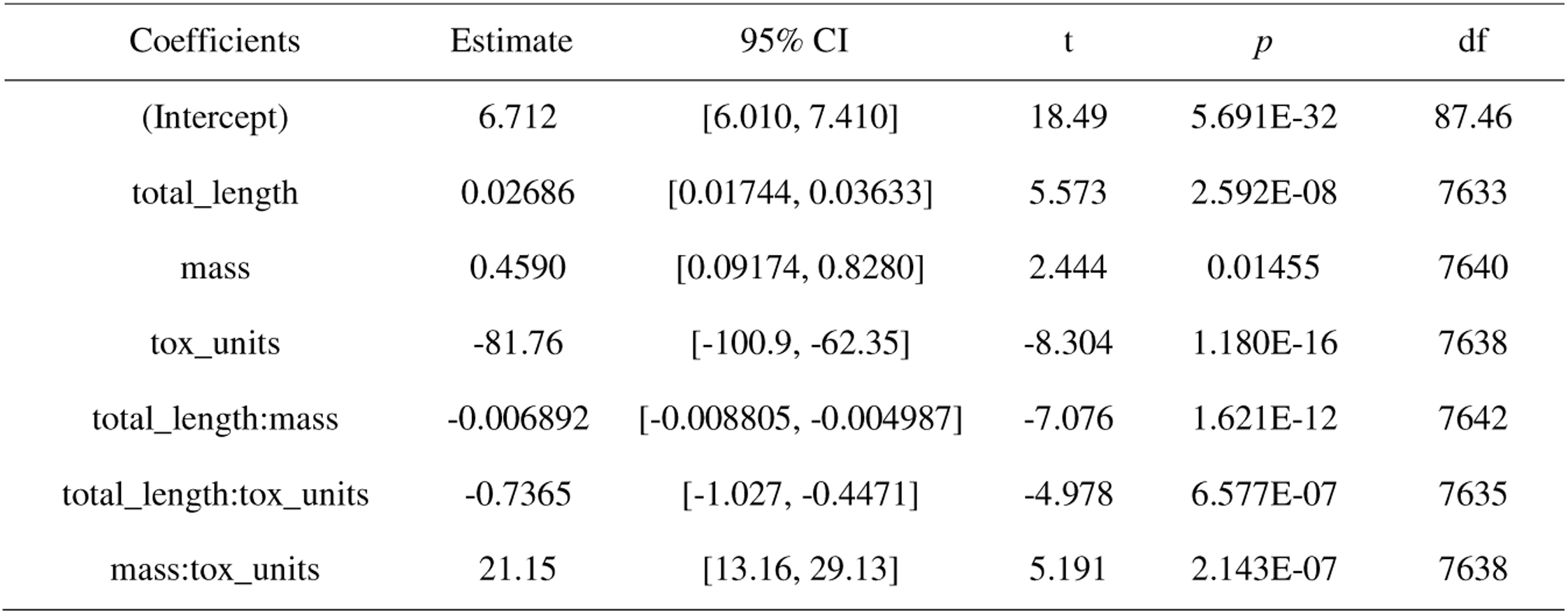
Linear mixed effect model results for Model A (reported in text). Model A is a linear mixed effect model evaluating the contribution of total length (mm), mass (g), toxicity units, and their interactions to survival time (hour). Experiment, tank, and bag are a nested random variable. Coefficient Estimate, 95% Confidence Interval, t value, p-value, and degrees of freedom (df) are from the model summary in R.

### Effects of ethanol storage on body size

HBBS and ARL larvae both declined in length and mass over the 120 day measurement period (ESM **Fig. S3**). For example, the total length of HBBS larvae (n = 110) initially ranged from 63.5 to 127.5 mm (μ = 95.1 mm; ESM **Fig. S3A**). The last measured total length (day 120) of HBBS larvae ranged from 56.0 to 119.5 mm (μ = 88.6 mm; ESM **Fig. S3A**). The mass of HBBS larvae (n = 110) initially ranged from 0.43 to 2.72 g (μ = 1.27 g; ESM **Fig. S3B**). The last measured mass of HBBS larvae ranged from 0.185 to 1.906 g (μ = 0.83 g; ESM **Fig. S3B**).

The nonlinear least squares model generated using the combined sample set (n = 664; site was included as a fixed variable to account for differences in ethanol storage volume) yielded a total length decay estimate as 4.483 • 10^-4^ mm/day (ESM **Table S4**). The linear regression model suggested a strong, positive relationship between predicted values and true values, with an average change in predicted initial length of 1.035mm for each 1mm increase in observed length post-storage after accounting for the effects of ethanol (*p* value < 2•10^-16^; ESM **Table S5**). The model explained a very large proportion of the variability in the initial length (R^2^ = 0.69). We used our model estimates to calculate initial total length, mass, and condition for our large TFM exposure experiment larvae by dividing the measured value by e raised to the negative estimate, multiplied by the number of days the larvae were stored in ethanol and found nearly identical results (ESM **Fig. S5**).

## Discussion

We describe a significant relationships between the total length, mass, and their interaction of larval sea lamprey and their survival time in TFM. Larger larval sea lamprey exhibit increased survival time in TFM, highlighting that this variable, if accounted for during field treatments, could improve treatment success. To conduct TFM treatments with the highest larval sea lamprey mortality, TFM treatment crews may want to consider larval size within a treatment stream and adjust treatment duration accordingly. We also describe the predicted survival time for larvae at the 10th, 25th, 50th, 75th, and 90th total length and mass percentiles within our data (**Fig. 2**; ESM **Table S6**), which can be used to justify the duration of treatment in selected streams based on surveyed larval sizes. For instance, streams where the average larval mass is greater than our 90th percentile would require at least an additional 30 minutes of treatment compared to streams where the average larval size closer to our 10th percentile (**Figure 2**; ESM **Table S6**). An earlier set of models incorporated condition factor (CF) as a co-variate, which was calculated as previously outlined for Pacific sea lamprey (Lampman et al., 2016) as 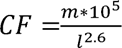 where *m* = mass (g) and *l* = length (mm). While, CF was not a significant contributor to reduced model AIC, perhaps because the equation is derived from a different species, the significant negative interaction between mass and total length reveals that condition is an important contributor to larval survival. Specifically, shorter and heavier larvae have longer survival times than longer and lighter-weight larvae (**Fig. 2**).

The observed correlation between sea lamprey size and survival time in TFM suggests the presence of a mechanism by which larger individuals can better withstand the toxic effects of the lampricide. Given that our linear mixed effect model including total length, mass, toxicity units, and their interactions explained a very large percentage of the variation in survival time (conditional R² = 0.8790), we suggest that the response of sea lamprey larvae to TFM is partly determined by larval size. While our primary focus was on describing the novel effects of larval size on survival to TFM exposure, we recognized that other variables—namely pH and TFM concentration —are well established as influential factors in TFM toxicity (Marking and Olsen 1975; Wilkie 2021). The inclusion of toxicity units in our model, despite our efforts to standardize experimental conditions, allowed us to account for known sources of variability that affect larval survival. By statistically controlling for these variables, we were able to more precisely isolate and evaluate the independent contributions of total length, mass, and their interaction on survival. This approach not only acknowledges the critical role of these environmental parameters in modulating TFM efficacy but also strengthens the inference that morphometric characteristics play a distinct and biologically relevant role in determining survival outcomes. We employed the LC90 target concentration of TFM to select individuals that exhibited prolonged survival, effectively sampling the right tail of the survival curve. This approach provided a robust sample size for future analysis of potential genetic factors associated with TFM survival.

In our study, the extended duration of survival for five individuals well beyond the typical treatment length of TFM in the field, that were neither the longest nor the heaviest larvae, indicate that some larvae may be capable of surviving treatment (**Fig. 3; ESM Fig. S4**). This observation is consistent with underlying variation in the population in terms of susceptibility to TFM and has implications for the potential for sea lamprey to evolve resistance to TFM, a subject of management interest (Christie et al., 2019; Dunlop et al., 2021). The development of resistance is a known consequence of repeated pesticide application (Délye et al., 2013; Georghiou and Taylor, 1977; Hawkins et al., 2019) and given that TFM has been used annually in freshwater streams in the Great Lakes Basin since 1958, resistance in sea lamprey is conceivable and an urgent concern (Christie et al., 2019; Dunlop et al., 2021). Prior research has suggested that a response to TFM-induced selection may already be underway in the Great Lakes: sea lamprey genomic and transcriptomic comparisons between treated and untreated populations of larval sea lamprey suggest that an adaptive, genetic response to TFM may drive incipient resistance (Yin et al., 2021). Sea lamprey could develop resistance to TFM through two broad mechanisms: (1) physiological adaptations or (2) evolution of their behavior or life history (Christie et al., 2019; Dunlop et al., 2018, 2021). Physiological resistance could involve the development of molecular mechanisms to more efficiently detoxify or neutralize TFM or to counteract the effects of the mitochondrial proton gradient; however, if larger individuals have a selective advantage for surviving treatment, and large size (or a correlated trait like growth rate) is heritable, then there is the potential for both physiological and life history resistance to TFM via selection for increases in body size. For example, larger sea lamprey could store additional glycogen, which would provide them with larger anaerobic energy reserves to generate ATP using anaerobic glycolysis. Such features could enable individuals to out-survive a typical TFM treatment. Although previous studies demonstrated no clear relationship between liver or muscle glycogen reserves and TFM tolerance, (Hlina et al. 2021; Muhametsafina et al., 2019), a recent study by Schueller et al. (2024) found a positive relationship between liver glycogen concentrations and the minimum lethal concentration (MLC = 12 h-LC^99.9^; Bills et al. 2003) of TFM to larval sea lamprey using stream-side toxicity tests. Larval sea lamprey also demonstrate a remarkable capacity to recover from TFM exposure when transferred to clean, TFM-free water, restoring energy stores such as glycogen and phosphocreatine within a few hours (Clifford et al. 2012). Collectively, these studies suggest that there may be a physiological advantage to being larger at the time of TFM exposure/treatments (Dunlop et al., 2021).

Contrary in part to our findings here, a handful of previous studies describe that energy stores and body size have little impact on TFM sensitivity (Hlina et al., 2021; Muhametsafina et al., 2019) and instead report that larvae exhibit differential TFM sensitivity based on season, which may be a result of the seasonally warmer waters enhancing their capacity to detoxify TFM (Hlina et al., 2021; Muhametsafina et al., 2019). Additionally, as pH and alkalinity of water increase, larvae exhibit increased tolerance as TFM changes its ionization state (reviewed in Wilkie et al., 2019), resulting in lower TFM uptake in more basic environments (Hlina et al., 2017). We accounted for pH within our analyses by proxy using toxicity units, controlled for alkalinity and water temperature during the experiment, and controlled for seasonality by using larvae collected within eight days in July to assess the contribution of total length and mass on survival time in TFM. Additional experiments may be needed to better weigh the contribution of other environmental and physiological components associated with differential survival in TFM (*e.g.*, temperature and alkalinity versus body mass and length).

In addition to quantifying the relationship between larval size and survival in TFM, we also characterized the relationship between larval body size and length after storage in ethanol. This information is valuable because future studies on resistance may involve preserving residual lamprey samples for subsequent analyses, and understanding the effects of ethanol storage on body length and mass will be essential. Similarly, accurate knowledge of preservation effects will benefit investigations into developmental milestones, such as size at metamorphosis, by informing decisions about sample storage methods. We provide an estimate of the decay rates of length and body mass, which should prove useful when back calculating the actual sizes of sea lamprey larvae prior to preservation (as demonstrated in ESM **Fig. S5**). By generating estimates for total length, mass, and condition, we evaluated the usefulness of each parameter to make accurate predictions of initial state values. The total length estimates generated from each sample set were of the same magnitude (on the scale of 10^-4^). Estimates from mass and condition were less accurate predictors of initial values, as reflected by smaller adjusted R^2^ values, when compared to those from total length. The best performing nonlinear regression model was derived from our combined sample set and provides a total length exponential decay estimate of 4.483•10^-4^ mm/day (ESM **Table S4**). Given that the model derived from our combined sample set had high predictability, our estimate may be useful across diverse larval sea lamprey collections. However, we also observed some variability in sample responses to ethanol preservation—while many larvae shrank over time, some exhibited minimal change or even increases in size (see ESM **Fig. S3**). These deviations from the expected dehydration pattern (i.e. exponentially shrinking in size) introduce a degree of uncertainty in back-calculations. Therefore, we advise researchers to use our correction equation with awareness of its limitations in cases where samples stored in ethanol may not follow typical shrinkage trends. Additionally, we observed much higher variability in our ARL data (ESM **Fig. S3B,F**) compared to the HBBS data (ESM **Fig. S3A,E**), likely due to different sample sizes and different combinations of researchers measuring each dataset, although many of the same researchers measured samples from each group of samples. Nevertheless, our most predictive model was that derived from the combined dataset of both groups of samples (ESM **Table S5**). Our model gives researchers additional options, apart from using formalin (Neave et al., 2006), for correcting for changes in body size after prolonged storage in 95% ethanol.

## Conclusion

Our study provides valuable insights into the relationship between larval sea lamprey size and survival time in TFM-treated environments. These findings suggest that duration and TFM dosage for routine TFM treatments within the Great Lakes basin could be improved if larval length and mass can be taken into consideration. Additionally, studies that store larval sea lamprey in 95% ethanol can now correct for storage time using a simple equation. We also emphasize the need for further research to better understand the underlying mechanisms between larger larval lamprey size and TFM survival to better inform sea lamprey management throughout the Great Lakes.

## Author contributions

Allison Nalesnik: Formal analysis, Investigation, Data curation, Writing-Original Draft, Writing-Review & Editing, Visualization, Supervision, Project administration. Emily Martin: Investigation, Data Curation, Writing-Review & Editing. Ian Kovacs: Investigation, Data Curation, Writing-Review & Editing. Connor Johnson: Investigation, Data curation, Writing-Review & Editing. Emma Carroll: Investigation, Data curation, Writing-Review & Editing. Aaron Jubar: Methodology, Resources, Writing-Review & Editing, Project administration. William Hemstrom: Formal analysis, Writing-Review & Editing. Michael Wilkie: Formal analysis, Writing-Review & Editing. Erin Dunlop: Conceptualization, Methodology, Writing-Review & Editing, Funding acquisition. Maria Sepulveda: Conceptualization, Methodology, Investigation, Resources, Writing-Review & Editing, Funding acquisition. Nicholas Johnson: Conceptualization, Methodology, Investigation, Resources, Data curation, Writing-Review & Editing, Supervision, Project administration, Funding acquisition. Mark Christie: Conceptualization, Methodology, Investigation, Resources, Writing-Review & Editing, Supervision, Project administration, Funding acquisition.

## Supporting information

SI Methods; Table S1; Table S2; Table S3; Table S4; Tabel S5; Table S6; Figure S1; Figure S2; Figure S3; Figure S4; Figure S5

## Acknowledgments

We thank the USFWS team responsible for collecting sea lamprey larvae for this experiment: Jordan Andrus, Philip Bennett, John Ewalt, Timothy Granger, Allen Keffer, David Keffer, Callie Kopp, Daniel McGarry, Lily Smalstig, Jenna Tews, and Matt Lipps. We also thank the USGS collection team and on-site personnel who provided hands-on assistance during the experiment: David Coburn, Tyler Bruning, Ed Benzer, Zachary Nordstrom, Meg Spens, and Brad Buechel. We thank Emma Vieregge and Karen Slaght for their support doing water quality analysis and range-finder assays. We thank Catherine Searle for her advice on our models. We also thank Robert Rode for his advice and assistance at the ARL, as well as undergraduate support from Lilly Gorney, Leah Kemple, and Grace Bowman. This research was funded by support to MRC, MSS, ESD, and NSJ from the Great Lakes Fishery Commission (grant number 2022_CHR_541010, 2022). Any use of trade, firm, or product names is for descriptive purposes only and does not imply endorsement by the U.S. Government.

## Conflict of interest

The authors declare that they have no conflict of interest.

## Data archiving

All code to reproduce the analyses reported in this manuscript are available via GitHub at https://github.com/ChristieLab/sea-lamprey-morphometrics. Data will be made available on Dryad.

## Notes

### Competing Interest Statement

The authors have declared no competing interest.

### Summary of Updates

Linear mixed effect model revised to include toxicity units and interactions with mass and total length; Removed one replicate (Exp1 tank S4) from analyses due to high variance in toxicity; Revised figures; Supplemental files updated

