## Supplementary material for "Larger larval sea lamprey (*Petromyzon marinus)* have longer survival times when exposed to the lampricide 3-trifluoromethyl-4-nitrophenol": SI Methods; Table S1; Table S2; Table S3; Table S4; Tabel S5; Table S6; Figure S1; Figure S2; Figure S3; Figure S4; Figure S5

**Supplementary Materials and Methods**

*Methods for repeated measurements of total length and mass of larvae stored in ethanol*

All measurement procedures took place in a fume hood. At the time of measuring, larvae were removed from their conical tube and briefly air dried under an air stream to remove excess surface ethanol. Larvae were then laid atop a ruler and total length (mm) was recorded. Next, larvae were placed on a weigh boat inside of a laboratory scale and mass (g) was recorded. Larvae were promptly returned to their original conical tube and placed aside. Larvae were measured one by one to prevent mistakenly returning a larva to the incorrect conical tube.

*Methods for measure of total length and mass of experiment larvae stored in ethanol*

Measurement procedures for the 8,611 larvae in our TFM exposure experiments were completed similarly to the procedure described above. Larvae that died during our TFM exposure experiments were stored together in containers ranging from 250mL to 1000mL. Each sample container housed only fish that died within the same hour, originating from the same tank and mesh bag. At the time of measuring, all larvae were poured into a plastic pitcher along with most of ethanol, avoiding excess desiccation from air exposure. Larvae were pulled from the pitcher one by one with tweezers and briefly air dried under an air stream to remove excess surface ethanol. Larvae were then laid atop a ruler and total length (mm) was recorded. Next, larvae were placed on a weigh boat inside of a laboratory scale and mass (g) was recorded. Larvae were returned to the original sample bottle with the original ethanol. All larvae from a given sample storage container were measured and returned to the same (original) container before another sample storage container was opened, proceeding one by one through sample containers.

**Acknowledgements**
Any use of trade, firm, or product names is for descriptive purposes only and does not imply endorsement by the U.S. Government.

**Supplemental Table 1.** Record of larval sea lamprey used in each 2022 experiment replicate.

| Replicate | Experiment Date | Bag ID | Total larvae | Collection | | Acclimation | |
| --- | --- | --- | --- | --- | --- | --- | --- |
|  |  |  |  | Date | Site | Arrived at HBBS | Holding Tank |
| 1 | 4 August 2022 | 1A | 72 | 12 July 2022 | Bridge St. Access | 13 July 2022 | 1 SMRT |
|  |  | 1B | 55 | 12 July 2022 | Bridge St. Access | 13 July 2022 | 1 SMRT |
|  |  | 1 | 147 | 19 July 2022 | Bridge St. Access | 21 July 2022 | 1 SMRT |
|  |  |  |  | 20 July 2022 | Bridge St. Access |  |  |
|  |  | 2 | 127 | 20 July 2022 | Bridge St. Access | 21 July 2022 | 1 SMRT |
|  |  | 3 | 137 | 20 July 2022 | Bridge St. Access | 21 July 2022 | 1 SMRT |
|  |  | 4 | 214 | 20 July 2022 | Bridge St. Access | 21 July 2022 | 1 SMRT |
|  |  |  |  | 20 July 2022 | Downstream Thornapple Rd. |  |  |
|  |  | 5 | 180 | 20 July 2022 | Downstream Thornapple Rd. | 22 July 2022 | 2 SMRT |
|  |  |  |  | 21 July 2022 | Bridge St. / Thornapple |  |  |
|  |  | 6 | 100 | 21 July 2022 | Bridge St. / Thornapple | 22 July 2022 | 2 SMRT |
|  |  | 7 | 115 | 21 July 2022 | Bridge St. / Thornapple | 22 July 2022 | 2 SMRT |
|  |  | 8 | 115 | 21 July 2022 | Bridge St. / Thornapple | 22 July 2022 | 2 SMRT |
|  |  | 9 | 247 | 21 July 2022 | Bridge St. / Thornapple | 22 July 2022 | 2 SMRT |
|  |  |  |  | 21 July 2022 | Downstream Thornapple Rd. |  |  |
|  |  | 10 | 190 | 21 July 2022 | Downstream Thornapple Rd. | 22 July 2022 | 2 SMRT |
|  |  |  |  | 21 July 2022 | Bridge St. Access |  |  |
| 2 | 6 August 2022 | 11 | 102 | 21 July 2022 | Bridge St. Access | 23 July 2022 | 3 SMRT |
|  |  |  |  | 22 July 2022 | Anderson Flat |  |  |
|  |  | 12 | 216 | 22 July 2022 | Anderson Flat | 23 July 2022 | 3 SMRT |
|  |  | 13 | 140 | 22 July 2022 | Anderson Flat | 23 July 2022 | 3 SMRT |
|  |  |  |  | 22 July 2022 | Devil’s Hole |  |  |
|  |  | 14 | 207 | 22 July 2022 | Devil’s Hole | 23 July 2022 | 3 SMRT |
|  |  | 15 | 205 | 22 July 2022 | Devil’s Hole | 23 July 2022 | 3 SMRT |
|  |  | 16 | 166 | 23 July 2022 | Devil’s Hole | 24 July 2022 | 3 SMRT |
|  |  |  |  | 23 July 2022 | Bridge St. Access |  |  |

**Supplemental Table 1.** Continued from the previous page.

| Replicate | Experiment Date | Bag ID | Total larvae | Collection | | Acclimation | |
| --- | --- | --- | --- | --- | --- | --- | --- |
|  |  |  |  | Date | Site | Arrived at HBBS | Holding Tank |
| 2 | 6 August 2022 | 17 | 127 | 23 July 2022 | Bridge St. Access | 24 July 2022 | 4 SMRT |
|  |  | 18 | 227 | 23 July 2022 | Bridge St. Access | 24 July 2022 | 4 SMRT |
|  |  | 19 | 248 | 23 July 2022 | Bridge St. Access | 24 July 2022 | 4 SMRT |
|  |  |  |  | 23 July 2022 | Devil’s Hole |  |  |
|  |  | 20 | 179 | 23 July 2022 | Devil’s Hole | 24 July 2022 | 4 SMRT |
|  |  | 21 | 91 | 23 July 2022 | Devil’s Hole | 24 July 2022 | 4 SMRT |
|  |  | 22 | 237 | 23 July 2022 | Devil’s Hole | 25 July 2022 | 4 SMRT |
|  |  |  |  | 24 July 2022 | Devil’s Hole |  |  |
| 3 | 9 August 2022 | 23 | 237 | 24 July2022 | Devil’s Hole | 25 July 2022 | 5 SMRT |
|  |  |  |  | 24 July2022 | Bridge St. Access |  |  |
|  |  | 24 | 153 | 24 July2022 | Bridge St. Access | 26 July 2022 | 5 SMRT |
|  |  |  |  | 25 July2022 | Bridge St. Access |  |  |
|  |  | 25 | 152 | 25 July2022 | Bridge St. Access | 26 July 2022 | 5 SMRT |
|  |  | 26 | 186 | 25 July 2022 | Devil’s Hole | 26 July 2022 | 5 SMRT |
|  |  | 27 | 223 | 25 July 2022 | Bridge St. Access | 27 July 2022 | 5 SMRT |
|  |  |  |  | 25 July 2022 | Devil’s Hole/ Thornapple |  |  |
|  |  |  |  | 25 July 2022 | Unspecified |  |  |
|  |  | 28 | 182 | 26 July 2022 | Bridge St. Access | 27 July 2022 | 5 SMRT |
|  |  | 29 | 239 | 26 July 2022 | Devil’s Hole | 27 July 2022 | 6 SMRT |
|  |  | 30 | 225 | 26 July 2022 | Bridge St. Access | 27 July 2022 | 6 SMRT |
|  |  | 31 | 161 | 26 July 2022 | Devil’s Hole | 27 July 2022 | 6 SMRT |
|  |  | 32 | 133 | 26 July 2022 | Devil’s Hole | 27 July 2022 | 6 SMRT |
|  |  | 33 | 166 | 26 July 2022 | Bridge St. Access | 27 July 2022 | 6 SMRT |
|  |  |  |  | 26 July 2022 | Devil’s Hole |  |  |
|  |  | 34 | 220 | 26 July 2022 | Devil’s Hole | 28 July 2022 | 6 SMRT |
|  |  |  |  | 27 July 2022 | Devil’s Hole |  |  |

**Supplemental Table 1.** Continued from the previous page.

| Replicate | Experiment Date | Bag ID | Total larvae | Collection | | Acclimation | |
| --- | --- | --- | --- | --- | --- | --- | --- |
|  |  |  |  | Date | Site | Arrived at HBBS | Holding Tank |
| 4 | 11 August 2022 | 35 | 212 | 27 July 2022 | Devil’s Hole | 28 July 22 | 7 SMRT |
|  |  | 36 | 179 | 27 July 2022 | Devil’s Hole | 28 July 22 | 7 SMRT |
|  |  | 37 | 223 | 27 July 2022 | Devil’s Hole | 28 July 22 | 7 SMRT |
|  |  | 38 | 169 | 27 July 2022 | Devil’s Hole | 28 July 22 | 7 SMRT |
|  |  | 39 | 178 | 27 July 2022 | Devil’s Hole | 28 July 22 | 7 SMRT |
|  |  | 40 | 189 | 27 July 2022 | Devil’s Hole | 28 July 22 | 7 SMRT |
|  |  | 41 | 228 | 27 July 2022 | Devil’s Hole | 28 July 22 | 8 SMRT |
|  |  | 42 | 139 | 28 July 2022 | Devil’s Hole | 28 July 22 | 8 SMRT |
|  |  | 43 | 244 | 25-26 July 2022 | Various locations | 26-27 July 2022 | 8 SMRT |
|  |  | 44 | 242 | 25-26 July 2022 | Various locations | 26-27 July 2022 | 8 SMRT |
|  |  | 45 | 205 | 25-26 July 2022 | Various locations | 26-27 July 2022 | 8 SMRT |
|  |  | 46 | 282 | 25-26 July 2022 | Various locations | 26-27 July 2022 | 8 SMRT |
| **Site Description (USFWS Site Description in SLMP) (Latitude, Longitude)** Bridge Street Access (Muskegon River at Sarell St. Public Access Site) (43.41517, -85.81133)  Thornapple Road (Muskegon River at High Rollway PFS (Thornapple PFS)) (43.41454667, -85.71821)  Anderson Flat (Muskegon River at Felch Ave. (Anderson Flats PFS)) (43.38885167, -85.82868333)  Devil’s Hole (Muskegon River at Schumaker Bayou (Devil’s Hole)) (43.42039, -85.73748) | | | | | | | |

**Supplemental Table 2.** Abiotic measurements across 2022 experimental replicates. Alkalinity and hardness are reported as milligrams/L calcium carbonate (CaCO_3_) and conductivity is reported as microsiemens per centimeter (µS/cm). Dissolved oxygen (DO) is reported as mg/L. Two 3-trifluoromethyl-4-nitrophenol (TFM) concentration measurements were taken: ‘inside’ refers to the tank center above the aeration stone and ‘outside’ refers to the outer edge of the tank.

| Replicate | Date | Tank | Alkalinity | Hardness | Conductivity | Hour | DO | pH | Temperature (°C) | | TFM Concentration (mg/L) | | |
| --- | --- | --- | --- | --- | --- | --- | --- | --- | --- | --- | --- | --- | --- |
|  |  |  |  |  |  |  |  |  | Air | Water | Mean | Inside | Outside |
| 1 | 4 August 2022 | S4 | 85 | 100 | 215 | 0 | 9.4 | 8.35 | 20.6 | 18.0 | 2.645 | 2.660 | 2.629 |
|  |  |  |  |  |  | 1 | 7.9 | 8.50 | 20.1 | 17.7 | 2.687 | 2.613 | 2.760 |
|  |  |  |  |  |  | 2 | 8.7 | 7.88 | 20.8 | 17.9 | 2.600 | 2.620 | 2.580 |
|  |  |  |  |  |  | 3 | 8.3 | 7.87 | 18.2 | 17.1 | 2.632 | 2.557 | 2.706 |
|  |  |  |  |  |  | 4 | 8.5 | 7.73 | 18.9 | 17.9 | 2.539 | 2.539 | 2.538 |
|  |  |  |  |  |  | 5 | 8.5 | 7.20 | 20.2 | 18.0 | 2.586 | 2.578 | 2.593 |
|  |  |  |  |  |  | 6 | 9.0 | 7.00 | 20.5 | 18.1 | 2.593 | 2.578 | 2.608 |
|  |  | S6 | 84 | 100 | 215 | 0 | 9.6 | 8.44 | 20.6 | 17.7 | 2.650 | 2.663 | 2.636 |
|  |  |  |  |  |  | 1 | 7.8 | 8.60 | 20.1 | 17.7 | 2.580 | 2.577 | 2.583 |
|  |  |  |  |  |  | 2 | 8.6 | 8.07 | 20.8 | 18.1 | 2.581 | 2.570 | 2.591 |
|  |  |  |  |  |  | 3 | 8.7 | 7.82 | 18.2 | 17.8 | 2.654 | 2.619 | 2.688 |
|  |  |  |  |  |  | 4 | 8.5 | 7.85 | 18.9 | 18.1 | 2.667 | 2.762 | 2.571 |
|  |  |  |  |  |  | 5 | 8.4 | 7.94 | 20.2 | 18.0 | 2.594 | 2.6 | 2.588 |
|  |  |  |  |  |  | 6 | 8.9 | 8.13 | 20.5 | 18.0 | 2.600 | 2.616 | 2.583 |
|  |  |  |  |  |  | 7 | 8.7 | 8.18 | 21.5 | 18.2 | 2.673 | 2.596 | 2.750 |
|  |  |  |  |  |  | 8 | 8.7 | 8.06 | 22.2 | 18.1 | 2.568 | 2.561 | 2.575 |
| 2 | 6 August 2022 | S4 | 86 | 100 | 211 | 0 | 9.5 | 8.47 | 20.9 | 17.7 | 2.743 | 2.702 | 2.784 |
|  |  |  |  |  |  | 1 | 8.5 | 8.05 | 21.3 | 17.8 | 2.693 | 2.751 | 2.634 |
|  |  |  |  |  |  | 2 | 8.1 | 7.86 | 22.1 | 18.2 | 2.539 | 2.611 | 2.467 |
|  |  |  |  |  |  | 3 | 7.6 | 7.79 | 22.7 | 18.1 | 2.574 | 2.565 | 2.583 |
|  |  |  |  |  |  | 4 | 7.9 | 7.84 | 23.2 | 18.3 | 2.572 | 2.574 | 2.570 |
|  |  |  |  |  |  | 5 | 8.4 | 7.94 | 23.7 | 18.7 | 2.584 | 2.591 | 2.577 |

**Supplemental Table 2.** Continued from the previous page.

| Replicate | Date | Tank | Alkalinity | Hardness | Conductivity | Hour | DO | pH | Temperature (°C) | | TFM Concentration (mg/L) | | |
| --- | --- | --- | --- | --- | --- | --- | --- | --- | --- | --- | --- | --- | --- |
|  |  |  |  |  |  |  |  |  | Air | Water | Mean | Inside | Outside |
| 2 | 6 August 2022 | S6 | 86 | 100 | 214 | 0 | 9.7 | 8.31 | 20.9 | 17.7 | 2.559 | 2.547 | 2.571 |
|  |  |  |  |  |  | 1 | 8.5 | 8.02 | 21.3 | 17.7 | 2.518 | 2.525 | 2.511 |
|  |  |  |  |  |  | 2 | 8.3 | 8.00 | 22.1 | 18.3 | 2.476 | 2.487 | 2.464 |
|  |  |  |  |  |  | 3 | 7.6 | 7.80 | 22.7 | 18.1 | 2.491 | 2.517 | 2.464 |
|  |  |  |  |  |  | 4 | 7.9 | 7.84 | 23.2 | 18.3 | 2.455 | 2.464 | 2.445 |
|  |  |  |  |  |  | 5 | 8.7 | 7.85 | 23.7 | 18.5 | 2.478 | 2.466 | 2.490 |
|  |  |  |  |  |  | 6 | 8.8 | 7.89 | 24.6 | 18.8 | 2.458 | 2.455 | 2.461 |
|  |  |  |  |  |  | 7 | 11.9 | 7.96 | 24.9 | 18.8 | 2.505 | 2.473 | 2.536 |
| 3 | 9 August 2022 | S4 | 86 | 101 | 215 | 0 | 9.2 | 8.22 | 16.5 | 16.7 | 2.589 | 2.634 | 2.544 |
|  |  |  |  |  |  | 1 | 9.4 | 8.08 | 17.9 | 16.6 | 2.528 | 2.525 | 2.531 |
|  |  |  |  |  |  | 2 | 8.8 | 8.05 | 17.9 | 16.8 | 2.488 | 2.477 | 2.498 |
|  |  |  |  |  |  | 3 | 8.8 | 7.92 | 18.0 | 16.8 | 2.540 | 2.510 | 2.569 |
|  |  |  |  |  |  | 4 | 8.9 | 7.98 | 18.7 | 16.7 | 2.501 | 2.503 | 2.499 |
|  |  |  |  |  |  | 5 | 9.1 | 7.98 | 18.7 | 16.7 | 2.498 | 2.495 | 2.501 |
|  |  |  |  |  |  | 6 | 9.4 | 8.12 | 20.0 | 16.9 | 2.497 | 2.496 | 2.497 |
|  |  |  |  |  |  | 7 | 9.4 | 8.22 | 20.6 | 17.0 | 2.485 | 2.482 | 2.487 |
|  |  |  |  |  |  | 8 | 9.4 | 8.25 | 21.4 | 17.1 | 2.563 | 2.563 | 2.563 |
|  |  |  |  |  |  | 9 | 9.4 | 8.30 | 21.7 | 17.2 | 2.520 | 2.507 | 2.533 |
|  |  |  |  |  |  | 10 | 9.5 | 8.39 | 22.3 | 17.1 | 2.558 | 2.555 | 2.560 |
|  |  |  |  |  |  | 11 | 9.5 | 8.49 | 22.4 | 17.3 | 2.550 | 2.551 | 2.548 |
|  |  |  |  |  |  | 12 | 9.5 | 8.37 | 22.3 | 17.3 | 2.565 | 2.576 | 2.553 |
|  |  |  |  |  |  | 13 | 9.4 | 8.41 | 22.3 | 17.4 | 2.593 | 2.585 | 2.601 |
|  |  |  |  |  |  | 14 | 9.4 | 8.38 | 21.6 | 17.3 | 2.498 | 2.522 | 2.474 |
|  |  |  |  |  |  | 15 | 9.5 | 8.30 | 22.2 | 17.5 | 2.534 | 2.523 | 2.545 |
|  |  |  |  |  |  | 16 | 9.6 | 8.36 | 22.3 | 17.6 | 2.521 | 2.532 | 2.509 |

**Supplemental Table 2.** Continued from the previous page.

| Replicate | Date | Tank | Alkalinity | Hardness | Conductivity | Hour | DO | pH | Temperature (°C) | | TFM Concentration (mg/L) | | |
| --- | --- | --- | --- | --- | --- | --- | --- | --- | --- | --- | --- | --- | --- |
|  |  |  |  |  |  |  |  |  | Air | Water | Mean | Inside | Outside |
| 3 | 9 August 2022 | S6 | 85 | 101 | 216 | 0 | 9.1 | 8.25 | 16.5 | 16.6 | 2.519 | 2.496 | 2.541 |
|  |  |  |  |  |  | 1 | 9.1 | 8.06 | 17.9 | 16.7 | 2.496 | 2.501 | 2.491 |
|  |  |  |  |  |  | 2 | 8.5 | 8.09 | 17.9 | 16.8 | 2.459 | 2.449 | 2.469 |
|  |  |  |  |  |  | 3 | 8.5 | 8.00 | 18.0 | 16.8 | 2.461 | 2.461 | 2.460 |
|  |  |  |  |  |  | 4 | 8.9 | 7.97 | 18.7 | 16.8 | 2.471 | 2.475 | 2.467 |
|  |  |  |  |  |  | 5 | 8.7 | 8.02 | 18.7 | 16.9 | 2.451 | 2.443 | 2.458 |
|  |  |  |  |  |  | 6 | 9.3 | 8.13 | 20.0 | 17.0 | 2.446 | 2.478 | 2.414 |
|  |  |  |  |  |  | 7 | 9.4 | 8.29 | 20.6 | 17.1 | 2.456 | 2.471 | 2.441 |
|  |  |  |  |  |  | 8 | 9.5 | 8.30 | 21.4 | 17.0 | 2.479 | 2.465 | 2.493 |
|  |  |  |  |  |  | 9 | 9.3 | 8.30 | 21.7 | 17.2 | 2.489 | 2.489 | 2.488 |
|  |  |  |  |  |  | 10 | 9.5 | 8.40 | 22.3 | 17.2 | 2.551 | 2.572 | 2.530 |
| 4 | 11 August 2022 | S4 | 86 | 100 | 210 | 0 | 9.1 | 8.04 | 19.8 | 17.6 | 2.503 | 2.508 | 2.498 |
|  |  |  |  |  |  | 1 | 8.1 | 8.01 | 18.7 | 17.6 | 2.420 | 2.415 | 2.425 |
|  |  |  |  |  |  | 2 | 8.1 | 8.03 | 18.9 | 17.4 | 2.463 | 2.463 | 2.463 |
|  |  |  |  |  |  | 3 | 8.7 | 7.91 | 18.9 | 17.8 | 2.541 | 2.546 | 2.536 |
|  |  |  |  |  |  | 4 | 8.2 | 7.96 | 19.8 | 17.4 | 2.418 | 2.433 | 2.402 |
|  |  |  |  |  |  | 5 | 9.1 | 7.96 | 19.5 | 17.4 | 2.458 | 2.456 | 2.460 |
|  |  |  |  |  |  | 6 | 9.4 | 8.14 | 20.1 | 17.4 | 2.431 | 2.433 | 2.429 |
|  |  |  |  |  |  | 7 | 8.6 | 8.24 | 20.5 | 17.5 | 2.480 | 2.479 | 2.480 |
|  |  |  |  |  |  | 8 | 9.8 | 8.19 | 20.7 | 17.5 | 2.484 | 2.475 | 2.492 |
|  |  |  |  |  |  | 9 | 9.8 | 8.25 | 21.0 | 17.5 | 2.482 | 2.479 | 2.485 |

**Supplemental Table 2.** Continued from the previous page.

| Replicate | Date | Tank | Alkalinity | Hardness | Conductivity | Hour | DO | pH | Temperature (°C) | | TFM Concentration (mg/L) | | |
| --- | --- | --- | --- | --- | --- | --- | --- | --- | --- | --- | --- | --- | --- |
|  |  |  |  |  |  |  |  |  | Air | Water | Mean | Inside | Outside |
| 4 | 11 August 2022 | S6 | 86 | 101 | 209 | 0 | 9.3 | 7.80 | 19.8 | 17.7 | 2.431 | 2.446 | 2.416 |
|  |  |  |  |  |  | 1 | 9.1 | 7.99 | 18.7 | 17.6 | 2.382 | 2.417 | 2.347 |
|  |  |  |  |  |  | 2 | 9.0 | 7.99 | 18.9 | 17.6 | 2.411 | 2.410 | 2.412 |
|  |  |  |  |  |  | 3 | 8.9 | 8.01 | 18.9 | 17.6 | 2.404 | 2.402 | 2.405 |
|  |  |  |  |  |  | 4 | 8.6 | 8.03 | 18.9 | 17.6 | 2.408 | 2.399 | 2.417 |
|  |  |  |  |  |  | 5 | 9.0 | 8.04 | 19.5 | 17.6 | 2.401 | 2.402 | 2.400 |
|  |  |  |  |  |  | 6 | 10.2 | 8.08 | 20.1 | 17.6 | 2.382 | 2.383 | 2.380 |
|  |  |  |  |  |  | 7 | 9.3 | 8.23 | 20.5 | 17.7 | 2.427 | 2.411 | 2.442 |
|  |  |  |  |  |  | 8 | 9.2 | 8.05 | 20.7 | 17.6 | 2.393 | 2.390 | 2.396 |
|  |  |  |  |  |  | 10 | 8.6 | 8.26 | 21.2 | 17.7 | 2.460 | 2.466 | 2.453 |
|  |  |  |  |  |  | 11 | 8.8 | 8.30 | 21.5 | 17.6 | 2.435 | 2.435 | 2.435 |
|  |  |  |  |  |  | 12 | 9.2 | 8.34 | 21.6 | 17.8 | 2.498 | 2.492 | 2.504 |
|  |  |  |  |  |  | 13 | 9.4 | 8.39 | 22.1 | 17.9 | 2.486 | 2.484 | 2.487 |
|  |  |  |  |  |  | 14 | 9.2 | 8.27 | 22.3 | 18.0 | 2.503 | 2.513 | 2.492 |
|  |  |  |  |  |  | 15 | 9.1 | 8.24 | 22.3 | 17.8 | 2.532 | 2.535 | 2.528 |
|  |  |  |  |  |  | 16 | 9.3 | 8.22 | 21.9 | 17.9 | 2.453 | 2.443 | 2.463 |
|  |  |  |  |  |  | 17 | 9.3 | 8.40 | 21.8 | 17.9 | 2.447 | 2.469 | 2.424 |
|  |  |  |  |  |  | 18 | 8.8 | 8.35 | 21.8 | 18.0 | 2.480 | 2.473 | 2.487 |

**Supplemental Table 3.** Extended linear mixed effect model results for Model A (reported in Table 2) and two other models (B-C) evaluating each size parameter with toxicity. Model A is a linear mixed effect model evaluating the contribution of total length, mass, the toxicity units, and their interactions to survival time (hour). All models have experiment, tank, and bag as a nested random variable. Coefficient Estimate, 95% Confidence Intervals, t value, p-value, and degrees of freedom (df) are from the model summary in R using 7,651 lamprey.

| **Coefficients** | **Estimate** | **95% CI** | **t** | ***p*** | **df** |
| --- | --- | --- | --- | --- | --- |
| Model A: hour ~ total_length*mass + total_length*tox_units + mass*tox_units + (1 \| experiment/tank/bag) | | | | | |
| (Intercept) | 6.712 | [6.010, 7.410] | 18.49 | 5.691E-32 | 87.46 |
| total_length | 0.02686 | [0.01744, 0.03633] | 5.573 | 2.592E-08 | 7633 |
| mass | 0.4590 | [0.09174, 0.8280] | 2.444 | 0.01455 | 7640 |
| tox_units | -81.76 | [-100.9, -62.35] | -8.304 | 1.180E-16 | 7638 |
| total_length:mass | -0.006892 | [-0.008805, -0.004987] | -7.076 | 1.621E-12 | 7642 |
| total_length:tox_units | -0.7365 | [-1.027, -0.4471] | -4.978 | 6.577E-07 | 7635 |
| mass:tox_units | 21.15 | [13.16, 29.13] | 5.191 | 2.143E-07 | 7638 |
| total_length | 0.02686 | [0.01744, 0.03633] | 5.573 | 2.592E-08 | 7633 |
| Model B: hour ~ total_length*tox_units + (1 \| experiment/tank/bag) | | | | | |
| (Intercept) | 7.915 | [7.415, 8.422] | 30.54 | 7.626E-16 | 16.34 |
| total_length | 0.01201 | [0.008167, 0.01592] | 6.082 | 1.242E-09 | 7642 |
| tox_units | -130.6 | [-141.12, -119.8] | -24.02 | 6.435E-123 | 7616 |
| total_length:tox_units | 0.02263 | [-0.08754, 0.1317] | 0.4047 | 0.6857 | 7635 |
| Model C: hour ~ mass*tox_units + (1 \| experiment/tank/bag) | | | | | |
| (Intercept) | 8.753 | [8.370, 9.146] | 47.32 | 1.786E-07 | 4.697 |
| mass | 0.1985 | [0.09722, 0.3013] | 3.818 | 0.0001358 | 7639 |
| tox_units | -133.0 | [-137.3, -128.5] | -59.15 | < 2.000E-16 | 7419 |
| mass:tox_units | 5.534 | [2.576, 8.466] | 3.685 | 0.0002307 | 7634 |

**Supplemental Table 4.** Results of nonlinear regression models describing the exponential decay of total length, mass, and condition (noted as VOI = Variable of Interest) as a function of ethanol storage time within each sample set: ARL (n=554), HBBS (n=110), and Combined (n=664). Each sample set consisted of independent data entries (one data entry per individual). For the nls model using the Combined sample set, site (either ARL or HBBS) was a fixed variable.

| Function: function(days_in_ethanol, VOI, B) {VOI / (exp(-B * days_in_ethanol))}  Model: nls(initial_VOI ~ expDecayModel(days_in_ethanol, VOI, B) | | | | | | | |
| --- | --- | --- | --- | --- | --- | --- | --- |
| **Sample Set** | **VOI** | **Estimate** | **Std. Error** | **t** | **p** | **df** | **Residual Std. Error** |
| ARL (n=554) | Total Length | 2.619e-04 | 3.451e-05 | 7.588 | 1.389e-13 | 553 | 3.545 |
|  | Mass | 0.007128 | 0.0001847 | 38.59 | 7.345e-159 | 552 | 0.222 |
|  | Condition | 0.007244 | 0.0002108 | 34.37 | 3.009e-139 | 552 | 0.3291 |
| HBBS (n=110) | Total Length | 9.298e-04 | 6.976e-05 | 13.33 | 1.251e-24 | 109 | 4.203 |
|  | Mass | 0.004813 | 0.0003033 | 15.87 | 4.278e-30 | 109 | .2605 |
|  | Condition | 0.003187 | 0.0002536 | 12.57 | 6.224e-23 | 109 | 0.1421 |
| Combined (n=664) | Total Length | 4.483e-04 | 3.222e-05 | 13.91 | 8.751e-39 | 663 | 3.810 |
|  | Mass | 0.006207 | 0.0001722 | 36.04 | 2.502e-158 | 663 | 0.2492 |
|  | Condition | 0.006621 | 0.0002072 | 31.95 | 2.560e-136 | 663 | 0.3258 |

**Supplemental Table 5.** Results of a linear model between known initial state values for total length, mass, and condition (measured at the time of storage on Day ‘0’ in Figure S2), and predicted initial state values after storage in ethanol generated with the model A estimates in Table S5. Individual estimates were generated by the nls model training on data without the data entry that was being predicted for. Each sample set consisted of independent data entries (one data entry per individual).

| **Sample Set** | **Variable** | **Estimate** | **Std. Error** | **t** | **p** | **df** | **Residual Std. Error** | **Adjusted R^2^** | **F** |
| --- | --- | --- | --- | --- | --- | --- | --- | --- | --- |
| ARL (n=554) | Total Length | 1.0134 | 0.01035 | 97.872 | <2e-16 | 552 | 3.411 | .9454 | 9579 on 1 and 552 df |
|  | Mass | 0.9143 | 0.01646 | 55.55 | 5.843e-228 | 551 | 0.1872 | 0.8482 | 3086 on 1 and 551 df |
|  | Condition | 0.8345 | 0.01684 | 49.55 | 3.746e-205 | 551 | 0.2708 | 0.8164 | 2455 on 1 and 551 df |
| HBBS (n=110) | Total Length | 0.9478 | 0.02765 | 34.284 | 7.186e-60 | 108 | 3.802 | 0.9151 | 1175 on 1 and 108 df |
|  | Mass | 0.8333 | 0.03498 | 23.82 | 8.354e-45 | 108 | 0.1984 | 0.8387 | 567.6 on 1 and 108 df |
|  | Condition | 0.5096 | 0.05350 | 9.524 | 5.631e-16 | 108 | 0.09825 | 0.4515 | 90.71 on 1 and 108 df |
| Combined (n=664) | Total Length | 1.0354 | 0.008545 | 125.167 | <2e-16 | 662 | 3.641 | 0.9568 | 1468 on 1 and 662 df |
|  | Mass | 0.8961 | 0.01434 | 62.51 | 6.501e-280 | 662 | 0.2016 | 0.8549 | 3907 on 1 and 662 df |
|  | Condition | 0.8219 | 0.01673 | 49.13 | 5.387e-223 | 662 | 0.2709 | 0.7844 | 2414 on 1 and 662 df |

**Supplemental Table 6.** Survival time (hours) predictions generated using estimates from Model A (see Table S4 for details) based on a larvae of length and weight at the 10th, 25th, 50th, 75th, and 90th percentile of our dataset of 7,651 larval sea lamprey. Mean toxicity units (0.03216) were used in the prediction model.

|  | | | **Mass (g) Percentiles** | | | | |
| --- | --- | --- | --- | --- | --- | --- | --- |
|  |  |  | 10% | 25% | 50% | 75% | 90% |
|  |  |  | 0.390 | 0.563 | 0.794 | 1.11 | 1.46 |
| **Total Length (mm) Percentiles** | 10% | 70.0 | 4.56 | 4.67 | 4.83 | 5.03 | 5.26 |
|  | 25% | 77.5 | 4.56 | 4.67 | 4.81 | 5.00 | 5.21 |
|  | 50% | 85.5 | 4.57 | 4.66 | 4.79 | 4.96 | 5.16 |
|  | 75% | 94.0 | 4.57 | 4.66 | 4.77 | 4.93 | 5.10 |
|  | 90% | 102.0 | 4.58 | 4.65 | 4.75 | 4.89 | 5.04 |

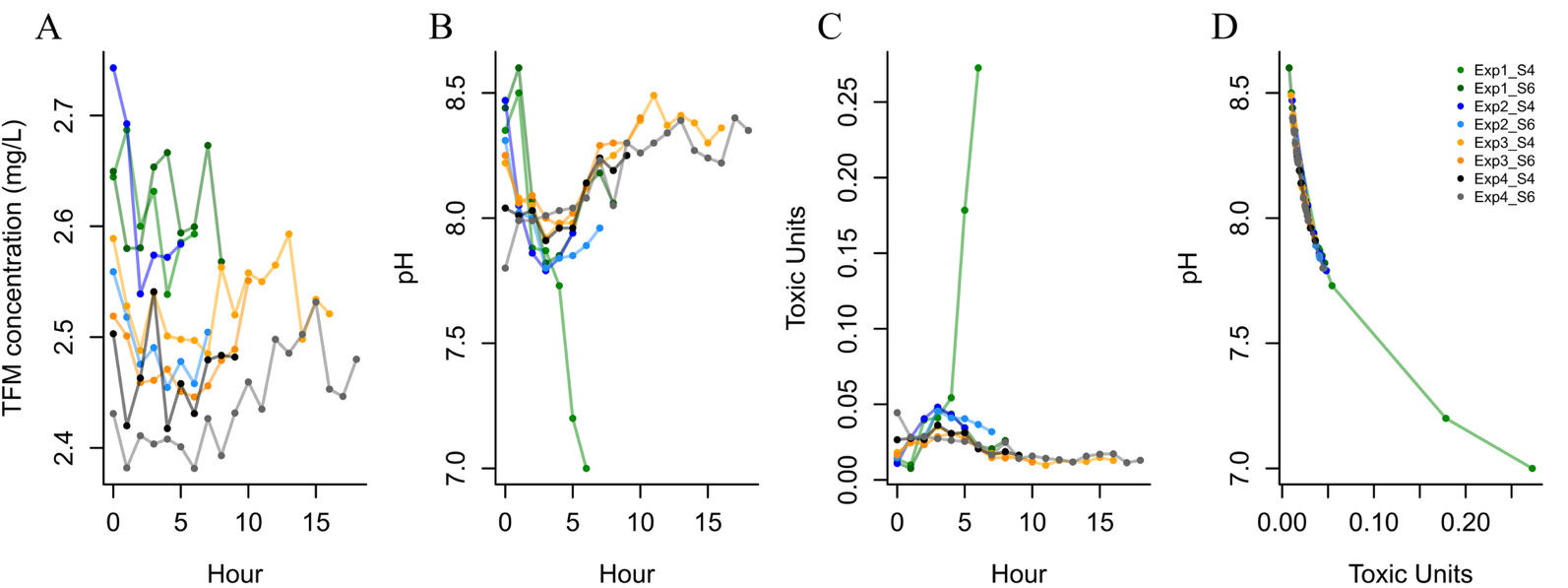
**Supplemental Figure 1. Changes in abiotic variables across experiment hours.** Fluctuations in (A) TFM concentration (mg/L), (B) pH, and (C) toxicity units over 18 hours. (D) The relationship between pH and toxicity units. Toxicity units are significantly correlated with survival time of larval sea lamprey in TFM (see Table S4 for model details).

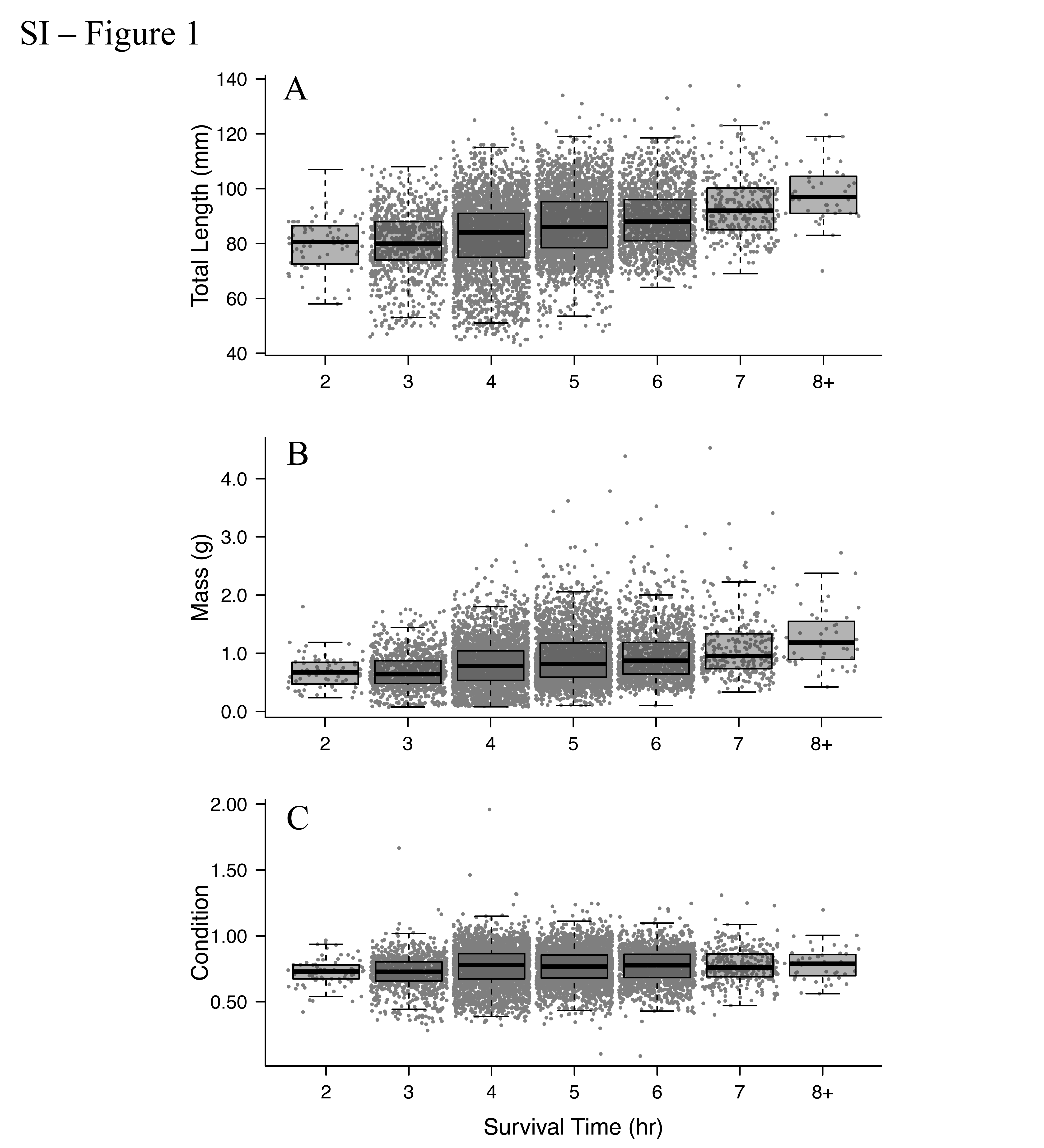

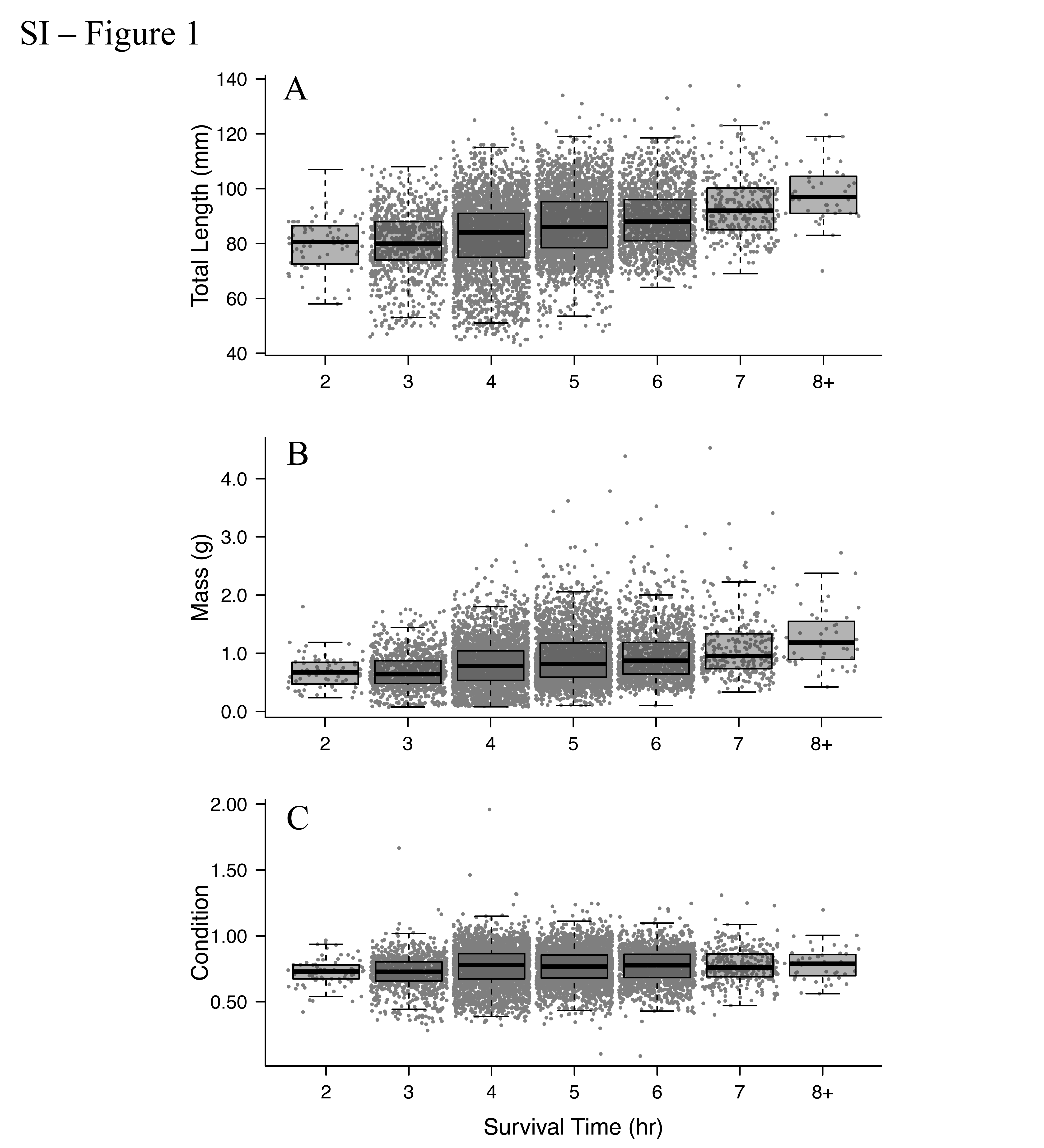

**Supplemental Figure 2. Relationship between morphometric variables and survival time in TFM illustrated for each experiment hour.** Relationships between (A) total length (mm) and (B) body mass (g) and survival of larval sea lamprey exposed to 9-h LC90 of 3-trifluoromethyl-4-nitrophenol (TFM; 2.3-2.4 mg/L) over up to 18 hours (n = 8,611 larvae). Larvae that survived beyond eight hours are binned within the group 8+ (See Figure S4 for survival for all hours). Data points are jittered 2.3 for visibility.

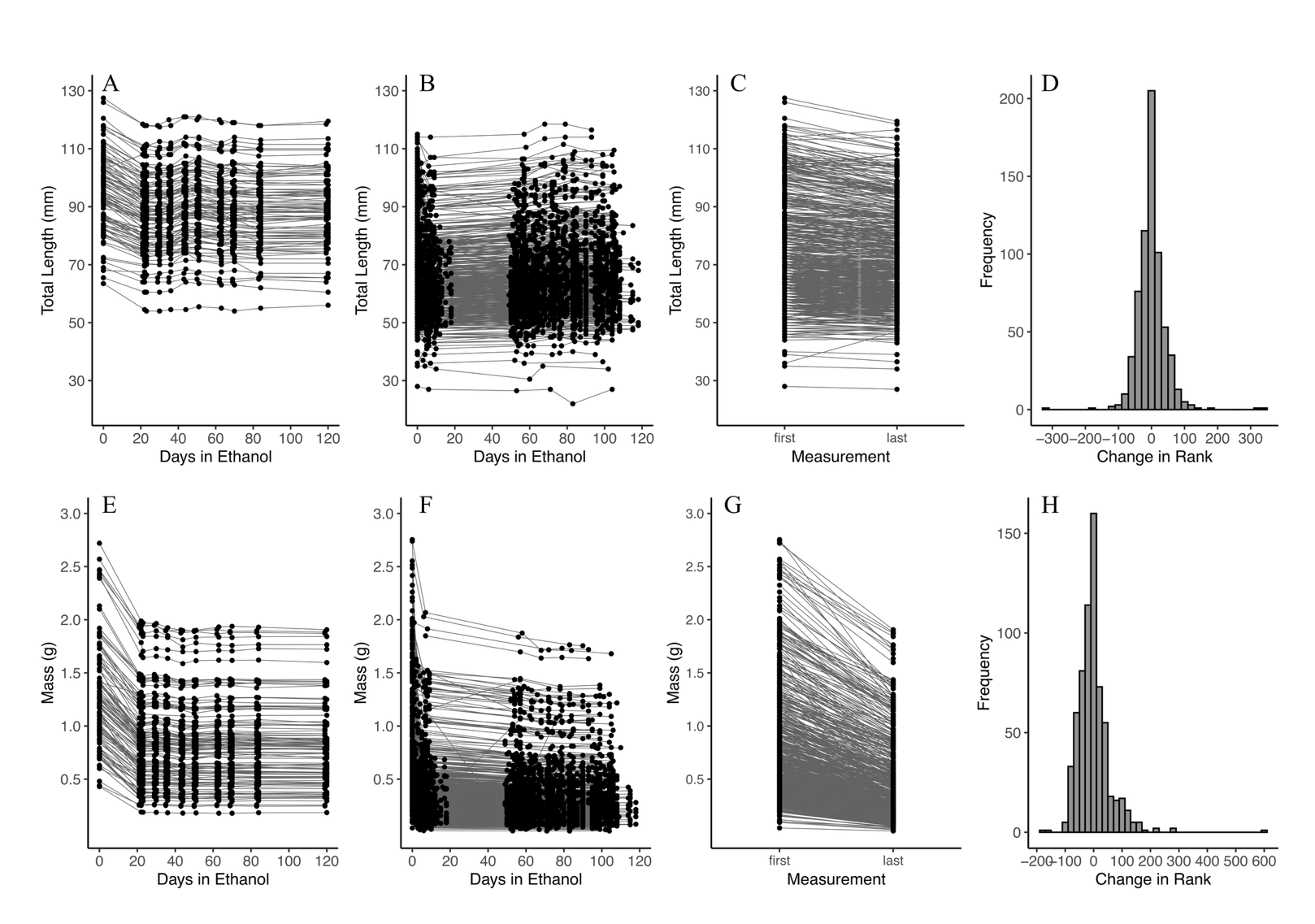
**Supplemental Figure 3. Changes in larvae total length and mass over time during storage in ethanol.** (A) Total length (mm) of larval sea lamprey from HBBS (n = 110) measured over successive days in ethanol storage. (B) Total length (mm) of larval sea lamprey from ARL (n = 554) measured over successive days in ethanol storage, initially ranging from 28 to 115 mm (μ = 68.1 mm) and then ranging from 27.0 to 116.5 mm (μ = 66.7 mm) at the last measurement. (C) Rank order of total length (mm) between first (time zero) and last (latest) measurements of larval sea lamprey stored in ethanol, with the shortest and longest larvae retaining their rank order. (D) Change in rank order position between first and last total length measurements, ranging from 0 to 345 (absolute μ rank change = 28.2). (E) Mass (g) of larval sea lamprey from HBBS (n = 110) measured over successive days in ethanol storage. (F) Mass (g) of larval sea lamprey from ARL (n = 554) measured over successive days in ethanol storage, initially ranging from 0.04 to 3.05 g (μ = 0.69 g) and then ranging from 0.01 to 1.76 g (μ = 0.34 g) at the last measurement. (G) Rank order of mass (g) between first and last measurements of ARL and HBBS larval sea lamprey stored in ethanol. (H) Absolute change in rank order position between first and last mass measurements for ARL and HBBS larvae, ranging from 0 to 598 (absolute μ rank change = 38.4).

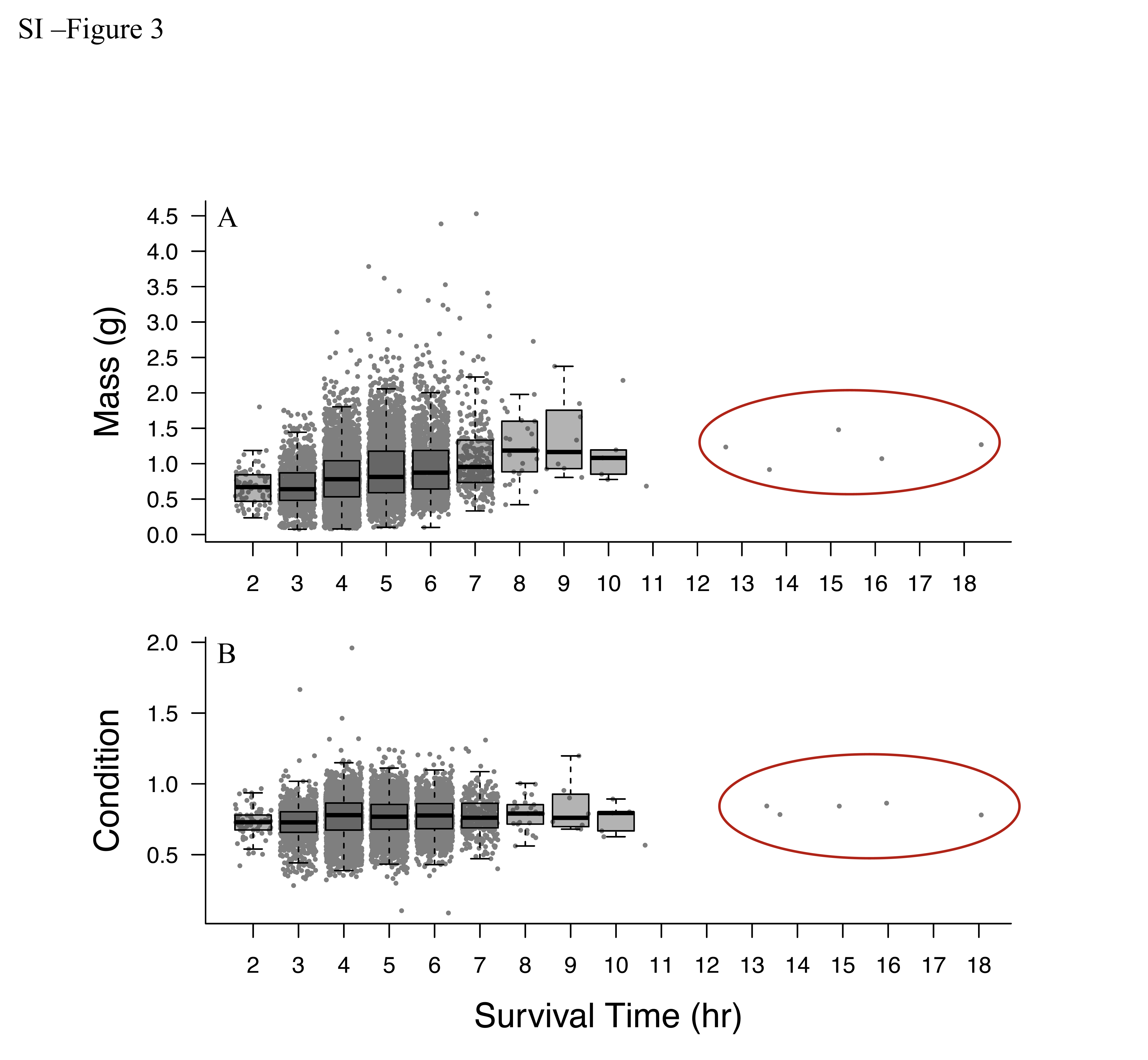

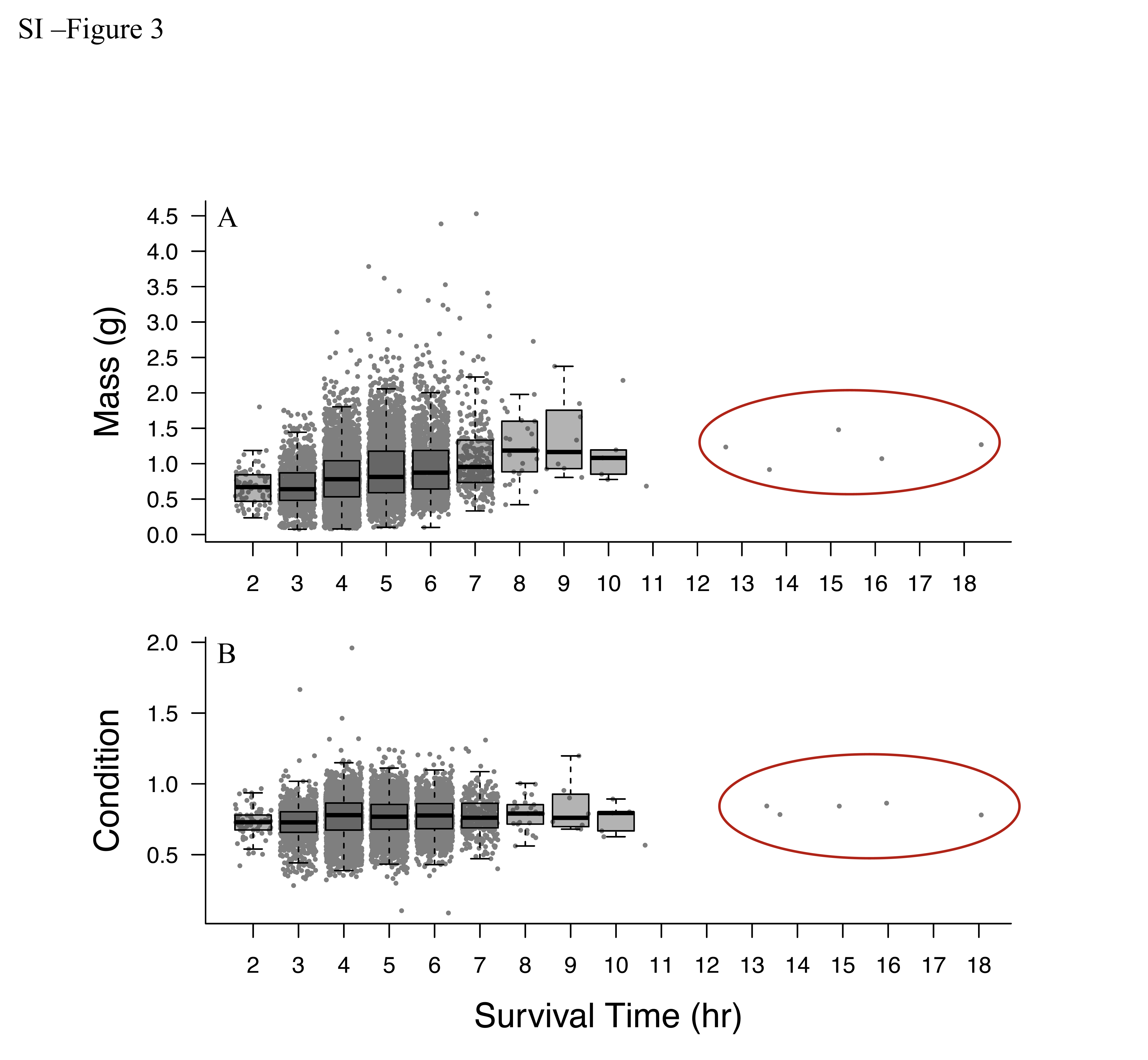
**Supplemental Figure 4. Relationship between mass and survival time in TFM.** Relationship between body mass (g) of larval sea lamprey exposed to 3-trifluoromethyl-4-nitrophenol (TFM; 2.3-2.4 mg L^-1^) for up to 18 hours (n = 8,611 larvae). Zero mortality occurred at hours one and 12. Five larval sea lamprey survived beyond 12 hours of TFM exposure (circled in red).

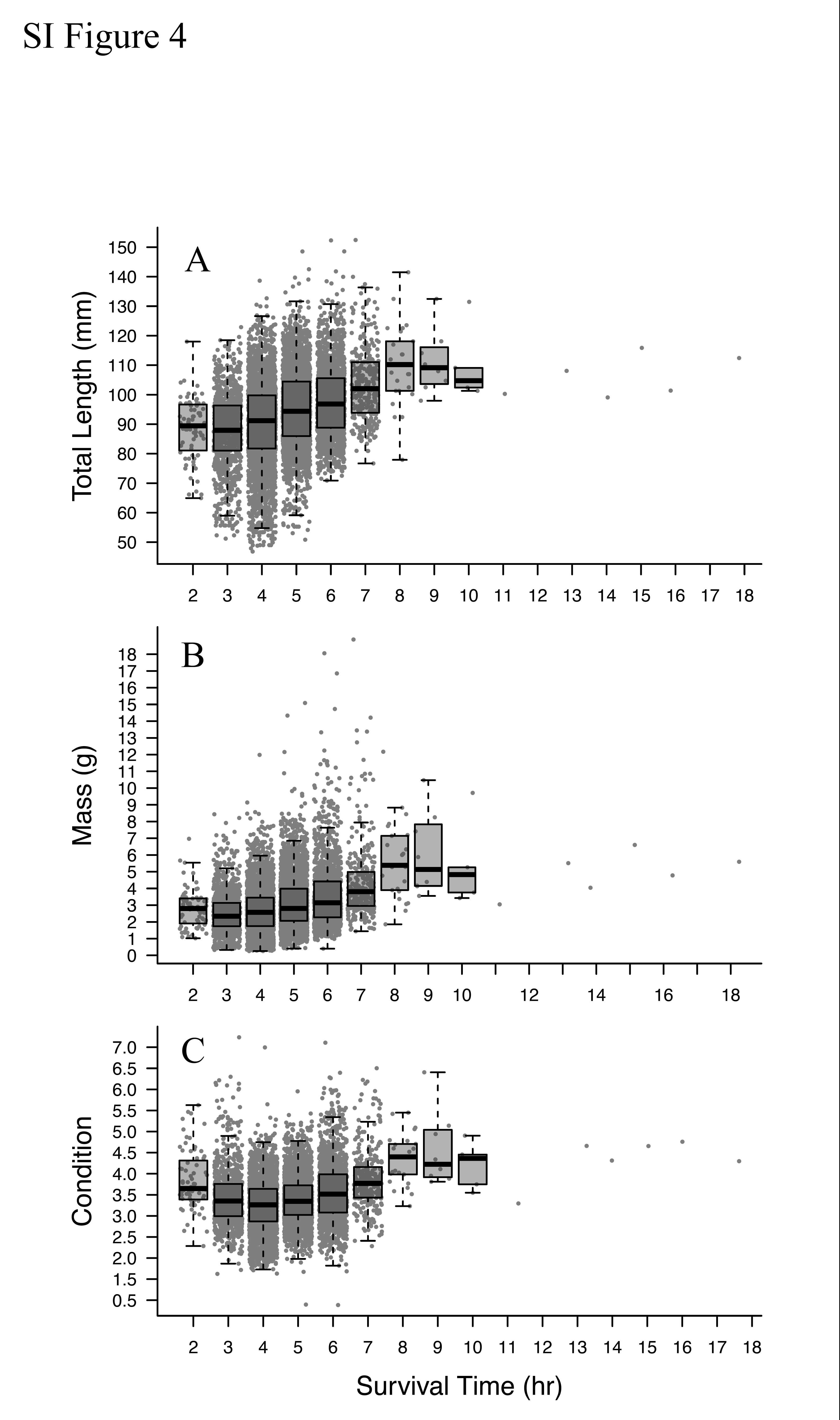

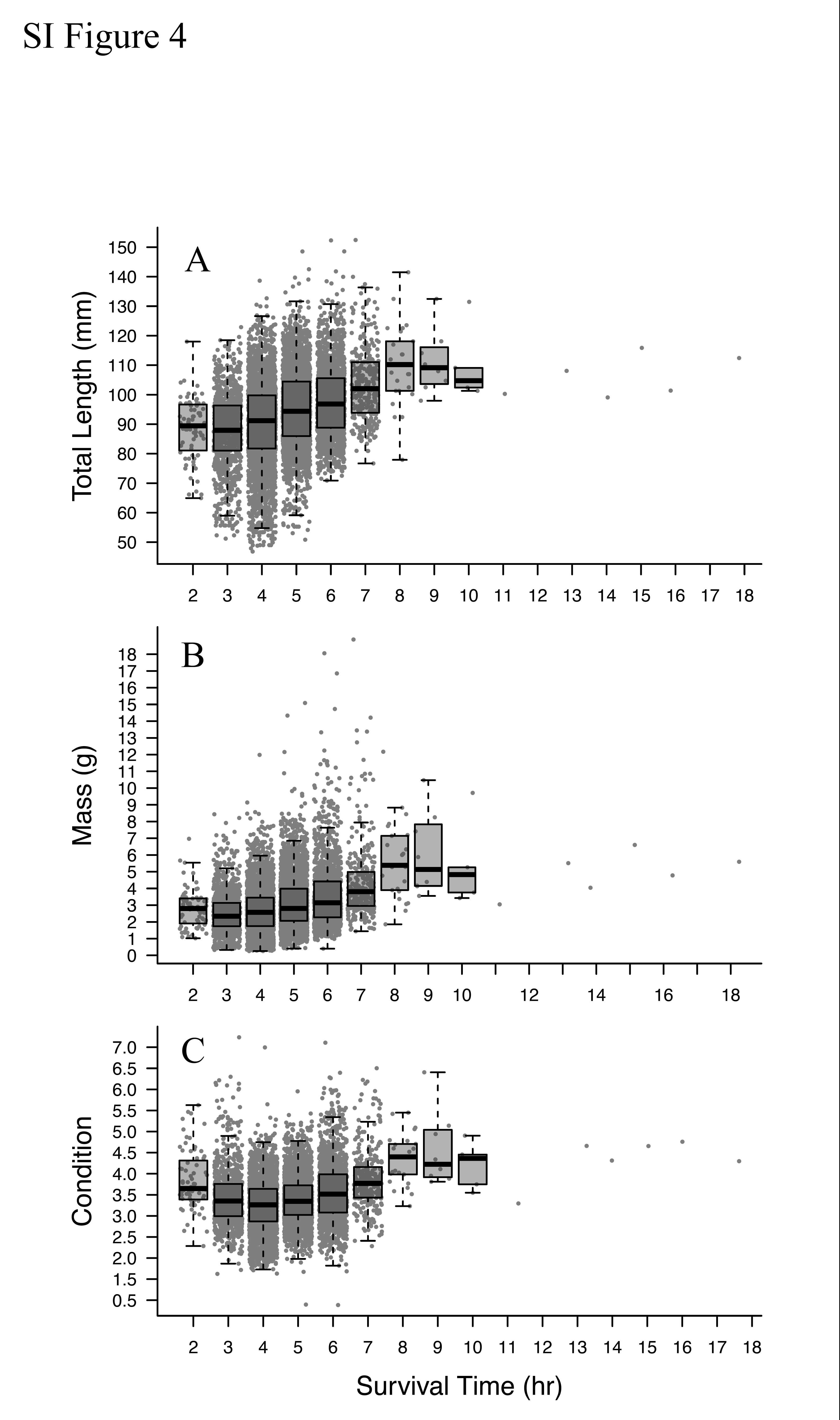

**Supplemental Figure 5. Adjusted morphometric variables after accounting for ethanol storage and survival time in TFM.** Relationships between (A) total length and (B) body mass and survival of larval sea lamprey exposed to 3-trifluoromethyl-4-nitrophenol (TFM; 2.52 ± 0.08 mg/L) up to 18 hours during(n = 8,611 larvae). Larvae size was adjusted based on decay rates estimated by our model: 4.483e-04 mm/day for total length and 0.006207 g/day for mass (Table S5).
